# Purinergic Ca^2+^ signaling as a novel mechanism of drug tolerance in BRAF mutant melanoma

**DOI:** 10.1101/2023.11.03.565532

**Authors:** Philip E. Stauffer, Jordon Brinkley, David Jacobson, Vito Quaranta, Darren R. Tyson

## Abstract

Drug tolerance is a major cause of relapse after cancer treatment. In spite of intensive efforts^1–9^, its molecular basis remains poorly understood, hampering actionable intervention. We report a previously unrecognized signaling mechanism supporting drug tolerance in BRAF-mutant melanoma treated with BRAF inhibitors that could be of general relevance to other cancers. Its key features are cell-intrinsic intracellular Ca^2+^ signaling initiated by P2X7 receptors (purinergic ligand-gated cation channels), and an enhanced ability for these Ca^2+^ signals to reactivate ERK1/2 in the drug-tolerant state. Extracellular ATP, virtually ubiquitous in living systems, is the ligand that can initiate Ca^2+^ spikes via P2X7 channels. ATP is abundant in the tumor microenvironment and is released by dying cells, ironically implicating treatment-initiated cancer cell death as a source of trophic stimuli that leads to ERK reactivation and drug tolerance. Such a mechanism immediately offers an explanation of the inevitable relapse after BRAFi treatment in BRAF-mutant melanoma, and points to actionable strategies to overcome it.

## INTRODUCTION

Drug tolerance, originally described in bacteria treated with antibiotics^10,11^, is recognized as a major challenge to cancer treatment, particularly with molecularly targeted agents. It is considered a major cause of residual disease, which may lead to tumor relapse and treatment failure^1,2,12–15^. Drug-tolerant cells survive treatment by non-genetic means, presumably via phenotypic adaptation^13,15–22^. Therefore, tolerance is distinct from resistance, which is based on pre-existing or acquired genetic mutations. The drug-tolerant state is thought to be reversible^3,13,21^, and it has been proposed that redirecting drug-tolerant cells into non-tolerant states across the phenotypic landscape should be possible^17,23^, such as by epigenetic modulators^21,24^. In spite of intensive efforts^1–9^, the molecular basis for drug tolerance remains poorly understood, hampering actionable intervention. An additional level of complexity to unravel is that drug tolerance has been described as a heterogeneous collection of distinct cell states^15,25–27^, making it challenging to effectively study and treat.

We previously described a distinct form of drug tolerance in BRAFi-treated BRAF-mutant melanoma that results in “idling” tumor cell populations^28,29^. After surviving initial drug exposure, drug-tolerant melanoma cells enter a state characterized by low, approximately equal rates of stochastic division and death events, leading to approximately zero net growth of the population; i.e., a cell population that idles, rather than being quiescent. In these idling cell populations, oncogenic BRAF stays inhibited^28^, but due to acquired drug tolerance ERK becomes reactivated and cell division still occurs sporadically. We proposed that these idling melanoma cell populations may be responsible for “residual disease” after BRAFi targeted therapy, which is thought to be a prelude to the acquisition of resistance-conferring mutations that ultimately defy treatment^15,30,31^.

Here, we report that, unexpectedly, purinergic signaling is a mechanism of drug tolerance in idling melanoma cell populations. Our data indicate that extracellular ATP (eATP), known to be abundant in the tumor microenvironment, triggers cytoplasmic Ca^2+^ ([Ca^2+^]_cyt_) spikes in BRAFi-adapted melanoma cells, through ATP-gated cation channels. We demonstrate that this purinergic signaling is then responsible for re-activation of ERK, which is known to promote viability and cell division under conditions of BRAFi. Further, we demonstrate that ERK in drug-tolerant cells is primed to respond to Ca^2+^ signals, illustrating the significance of these cell-intrinsic [Ca^2+^]_cyt_ spikes to the drug-tolerant state. To the best of our knowledge, purinergic signaling has not been described as a mechanism of ERK-reactivation in drug tolerance before and could produce actionable insights.

## RESULTS

### BRAFi induces cell-intrinsic Ca^2+^ signaling in drug-tolerant cells

Previous^32^ and more recent (Fig. 1A) findings by our lab implicated changes in ion channel expression and Ca^2+^ signaling as a mechanism of adaptation and drug tolerance to BRAFi in BRAF-mutant melanoma. Ion channels and Ca^2+^ signaling have been studied in the context of cellular physiology, and to a much lesser extent in cancer^33–38^, but no studies have directly explored their potential involvement in drug tolerance (see Discussion). To investigate this possibility, we explored changes in Ca^2+^ signaling in BRAFi-treated drug-tolerant idling populations (≥3 days BRAFi). By measuring changes in [Ca^2+^]_cyt_ using the Ca^2+^ dye, Fura-2-AM, we found that frequent spontaneous spikes of [Ca^2+^]_cyt_ were pronounced in drug-tolerant idling cells, occurring in up to 60% of the population (Fig. 1B, Fig. 1C, Supplementary movie M1, and Supplementary Fig. S1). Spike patterns from individual cell traces appeared in various forms, as represented in Fig. 1B. Cell traces varied with respect to the magnitude, shape, and frequency of spiking, ranging from near constant spiking to only one or two spikes during the imaging period (Fig. 1B and Supplementary Fig. S2). Within cell traces, complex signaling patterns emerged across the population, with some cells displaying multiple spike shapes, possibly hinting at the heterogeneity that underlies drug tolerance (see Discussion).

**Figure 1.**
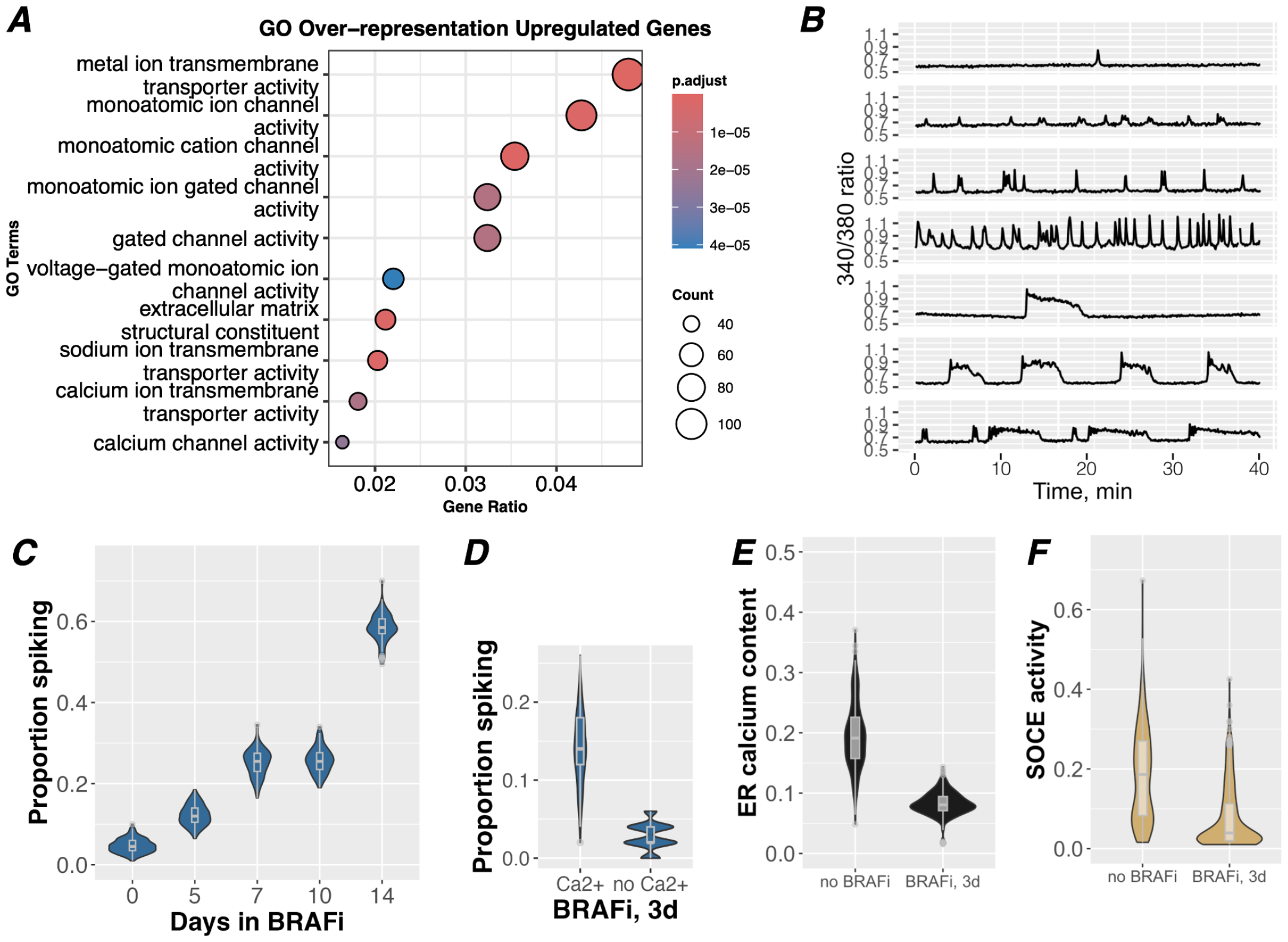
Ca^2+^ transport and cytoplasmic Ca^2+^ regulation are associated with drug tolerance. *A)* GO term representation of significantly upregulated genes from bulk RNAseq data across four melanoma cell lines under BRAFi conditions for 8 days, a treatment previously shown to induce an idling population of drug-tolerant cells^**28**,**32**^. Nine out of 10 of the top GO terms relate explicitly to ions and ion channels, two of which relate specifically to Ca^2+^, though Ca^2+^ related genes are encompassed in nearly all of these GO terms. *B)* Representative traces from imaging of [Ca^2+^]_cyt_ spikes with Fura-2 in A375 cells treated with 8 uM PLX4720 for 14 days. Data is plotted as a ratio of Fura-2 excitation/emission at 340/380 nm; An increase in the ratio indicates an increase in cytoplasmic Ca^2+^. A variety of spike patterns, magnitudes, and frequencies observed in the population are represented here. *C)* A375 cells were treated with 8 uM PLX4720 for the indicated number of days before imaging with Fura-2 for 40 minutes to detect [Ca^2+^]_cyt_ spikes. Traces were identified as having at least one spike or no spikes using an automated spike detection algorithm. Bootstrapping was performed iteratively to randomly select cells in the datasets to estimate proportions of the population that experience at least one spike. With increasing treatment time, there is a clear increase in the activity of [Ca^2+^]_cyt_ spikes. *D)* A375 cells treated with 8 μM PLX4720 for 3 days were imaged with Calbryte-520 to detect [Ca^2+^]_cyt_ spikes under conditions with and without e[Ca^2+^]. Traces were manually assessed for the presence of [Ca^2+^]_cyt_ spikes and proportions of spiking cells were determined with bootstrapping. *E)* Store operated Ca^2+^ entry (SOCE) assays were performed on A375 cells treated with 8 μM PLX4720, or vehicle, for 3 days to quantify ER Ca^2+^ content and activity levels of SOCE. Integrals were taken to quantify Ca^2+^ released during the ER Ca^2+^ phase, separate from the SOCE phase of the assay. Values within individual cells are background normalized and plotted to generate these distributions.

Spiking activity became more prevalent across the population with increasing BRAFi treatment times (Fig. 1C), demonstrating this signaling is a time-dependent response that coincides with acquisition of drug tolerance. In contrast, few to no spikes were detected in cells either in the absence of BRAFi or at 60 minutes BRAFi (referred to here as drug-sensitive populations)(Fig. 1C and Supplementary Fig. S3). Furthermore, when cells were treated with one of the clinical standard-of-care combinations of BRAFi (dabrafenib, 1 μM) and MEKi (trametinib, 100 nM) for 3 days, similar spontaneous [Ca^2+^]_cyt_ spikes were also observed (Supplementary Fig. S1), indicating this signaling mechanism may also be relevant to melanoma cell survival in clinical settings.

Identifying the sources of these [Ca^2+^]_cyt_ signals may elucidate their significance and molecular function in the drug-tolerant state. Broadly, [Ca^2+^]_cyt_ spikes may originate from intracellular or extracellular Ca^2+^ sources, or a combination of both. Ligand activation of plasma membrane receptors (i.e. G-protein-coupled receptors) may lead to [Ca^2+^]_cyt_ release from intracellular Ca^2+^ stores in the endoplasmic reticulum (ER), principally through inositol triphosphate (IP3) receptors (ITPRs)^39^. Alternatively, [Ca^2+^]_cyt_ spikes could be a result of influx of extracellular Ca^2+^ (e[Ca^2+^]) through plasma membrane ion channels, down an electrochemical gradient. To differentiate between these potential sources, cells were switched to a Ca^2+^ free buffer immediately before imaging. In the absence of e[Ca^2+^], spontaneous [Ca^2+^]_cyt_ spikes are diminished to approximately 2.5% of the idling population (Fig. 1D and Supplementary Fig. S4), demonstrating the importance of plasmalemma Ca^2+^ flux. Conversely, when we assessed the potential contribution of intracellular Ca^2+^ flux, we found that ER Ca^2+^ stores were substantially reduced in the drug-tolerant population (Fig. 1E and Supplementary Fig. S5), suggesting a diminished capacity for signaling from intracellular Ca^2+^ sources. Similarly, the mechanism for refilling ER stores upon depletion, i.e. as a result of intracellular Ca^2+^ signaling, via store operated Ca^2+^ entry (SOCE)^40^ was also substantially reduced (Fig. 1E). Taken together, these data suggest that drug-tolerant cells may have a blunted capacity to maintain Ca^2+^ signaling from intracellular sources and/or from SOCE activity itself^41^. Moreover, mRNA expression levels of ITPRs, which are central to ER Ca^2+^ signaling, are significantly reduced in drug-tolerant cells (Supplementary Fig. S6). These findings directed us to focus on a role of plasmalemmal flux of e[Ca^2+^] rather than release of intracellular Ca^2+^ stores as a primary mechanism of these [Ca^2+^]_cyt_ spikes.

### Purinergic Ca2+ signaling is a mechanism of ERK reactivation in drug-tolerant idling cells

To explore which ion channel(s) may be responsible for the abundance of plasmalemma Ca^2+^ flux, we visualized expression changes of specific genes in BRAF-mutant melanoma cell lines in the idling drug-tolerant state (8-days under BRAFi conditions). Consistent with the observed [Ca^2+^]_cyt_ spikes, the surviving cell population exhibited significant upregulation of numerous genes involved in Ca^2+^ signaling (Fig. 2A and Supplementary Figs. S7, S8). Notably, the ATP-gated, monoatomic cation conducting channel Purinergic Receptor P2X7 (gene name P2RX7) was one of the highest upregulated genes in the dataset, with a >250 fold increase at the mRNA level in drug-tolerant A375 cells and >6.6 fold across all drug-tolerant cell lines combined (Figs. 2A, 2B, and Supplementary Figs. S7, S8).

**Figure 2.**
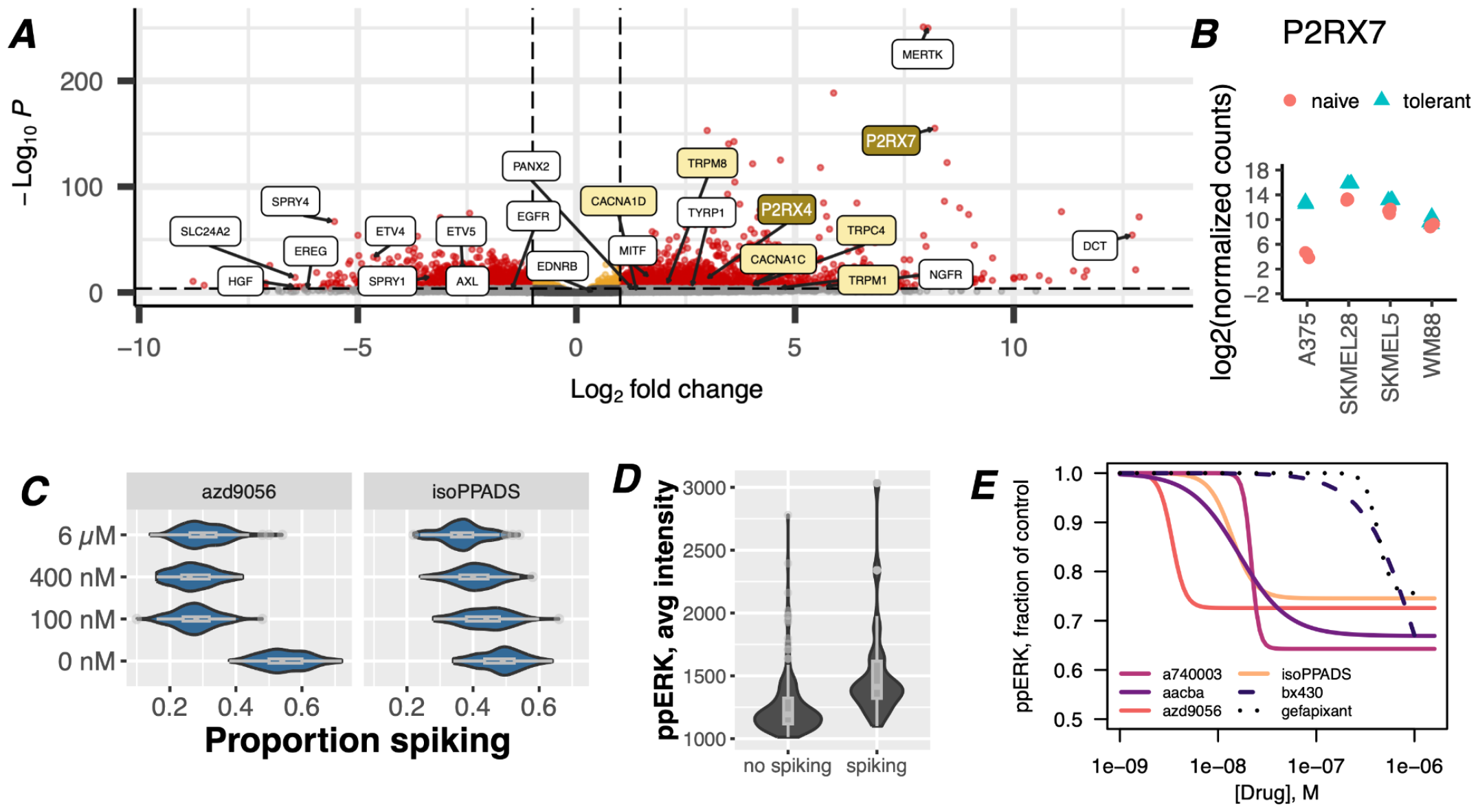
Upregulation and activation of purinergic receptors induce spontaneous cytoplasmic Ca^2+^ spiking and activation of ERK in drug-tolerant cells. *A)* Volcano plot of up- and down-regulated genes in melanoma cells under BRAFi conditions for 8 days. Data are from bulk RNAseq of a representative cell line, A375, out of four that were tested (all data combined are shown in Supplementary Figures S7 and S8). Reduced MEK-dependent genes and increased melanocyte differentiation genes changed as expected for continued inhibition of oncogenic BRAF activity^**42–44**^. Transcripts for the Ca^2+^ channel P2RX7 (upper right) are enriched by 250 fold, with a p value of < 10^−150^. *B)* Log2 fold change in expression of P2RX7 transcripts before and after BRAFi treatment (8 days) in four BRAF-mutant melanoma cell lines. *C)* A375 cells were treated with 8 uM PLX4720 for 3 days before being treated with the P2X7 inhibitors, AZD9056 and isoPPADS (or vehicle), immediately before imaging with Calbryte-520 to detect [Ca^2+^]_cyt_ spikes. Traces were manually analyzed for spiking activity. Bootstrapping was performed to generate proportions of spiking cells presented in the figure. These drugs had inhibitory effects on spiking activity at concentrations expected from their known IC50 values. *D)* A375 cells treated with 8 μM PLX4720 for 3 days were imaged with Calbryte-520 to detect [Ca^2+^]_cyt_ spikes, followed by fixation and staining for ppERK. Plotting ppERK staining intensity for cells classified as spiking or no spiking by manual assessment revealed higher ppERK staining in spiking cells. *E)* A375 cells treated with BRAFi for 3 days were incubated with a panel of P2X7 inhibitors (or vehicle) for one hour and assessed for levels of ppERK intensity at the single-cell level with and without inhibitors present. Cells staining positive for ppERK were quantified and calculated as ratios (fraction) of vehicle control. Log-logistic models of concentration dependent effects of each P2Xi were fit to the data and are represented by the lines as indicated.

P2X7 belongs to a family of ATP responsive ionotropic (P2X) and metabotropic (P2Y) purinergic receptors that have been studied extensively in the context of Ca^2+^ signaling in the nervous system and the immune system, largely with respect to cellular signaling, inflammation, and inflammatory pathologies^45–47^. However, purinergic signaling has been broadly overlooked in the context of oncogenic BRAF activity. Nonetheless, the Human Protein Atlas and The Cancer Genome Atlas report a trend for P2RX7 enrichment at both transcript and protein levels in skin and melanoma tissues, and worse clinical outcomes are associated with high expression (Supplementary Fig. S9)^48,49^. To specifically explore whether the spontaneous [Ca^2+^]_cyt_ spikes in drug-tolerant melanoma cells originate from P2X7 receptors, we adapted and optimized high-throughput equipment to image cells in the presence of unrelated P2X7 inhibitors (P2X7i). We found that approximately 24-50% fewer drug-tolerant cells experienced [Ca^2+^]_cyt_ spikes (Fig. 2C) in the presence of P2X7i, demonstrating that P2X7 signaling is a major mechanism of these BRAFi induced [Ca^2+^]_cyt_ spikes.

We then directed our attention to possible molecular mechanisms that may link P2X-derived [Ca^2+^]_cyt_ spikes to drug tolerance. ERK (re)activation is observed in and has been shown to be essential for drug tolerance because it is well known to promote cell survival and proliferation^50,51^. Ca^2+^ signals have been shown to activate the MAPK pathway in many cell types^52–62^, though it has never been considered in the context of drug tolerance. By performing single-cell Ca^2+^ measurements on drug-tolerant cells, followed by fixation and staining for phospho-ERK T202/Y204 (ppERK), we found that drug-tolerant cells with [Ca^2+^]_cyt_ spikes have significantly greater ppERK staining (Fig. 2D). Accordingly, co-treatment with P2X7 inhibitors reduced the proportion of cells with ERK reactivation (Fig. 2E), at concentrations below their *in vitro* IC50 values^63,64^, confirming purinergic [Ca^2+^]_cyt_ spikes are mitogenic. Drugs that target P2X4 (BX430) or P2X3 (gefapixant), also displayed activity, but only at concentrations significantly above their IC50 values, demonstrating the particular potency of P2X7 targeted inhibitors. Notably, approximately the same proportion of cells that lost [Ca^2+^]_cyt_ spikes also lost ppERK staining, as would be expected from a causal relationship.

### ATP stimulates Ca^2+^ signaling and ERK reactivation via P2X7 receptors in drug-tolerant idling cells

ATP, a physiological agonist of P2X receptors, is known to be abundant in the tumor microenvironment^36^. If ATP were to activate ppERK in the presence of BRAFi, an intriguing relationship to drug tolerance in melanoma tumors could emerge. Indeed, exogenous ATP induces ppERK activation in drug-tolerant, but not drug-sensitive cells (Fig. 3A), indicating relevance to an adaptive response. As expected, ATP stimulation of ppERK corresponds to activation of [Ca^2+^]_cyt_ spikes (Fig. 3B, ‘no ATP, 0’ vs ‘0’) that resemble the spontaneous [Ca^2+^]_cyt_ spikes observed in Fig. 1B (Supplementary Fig. S10), which occur in the absence of exogenous ATP. Inhibition of these ATP induced [Ca^2+^]_cyt_ spikes (Fig. 3B) results in the reduction of ATP stimulated ppERK staining (Fig. 3C). Additionally, ATP-induced cytoplasmic Ca^2+^ levels were sustained at higher levels in drug-tolerant cells than drug-sensitive cells, indicative of potentiation of ATP-induced effects in the drug-tolerant condition (Supplementary Fig. S11). These results demonstrate that ATP-induced Ca^2+^ signaling leads to increased ppERK in a P2X7 receptor-dependent manner, potentially enhanced by prolonged signaling dynamics.

**Figure 3.**
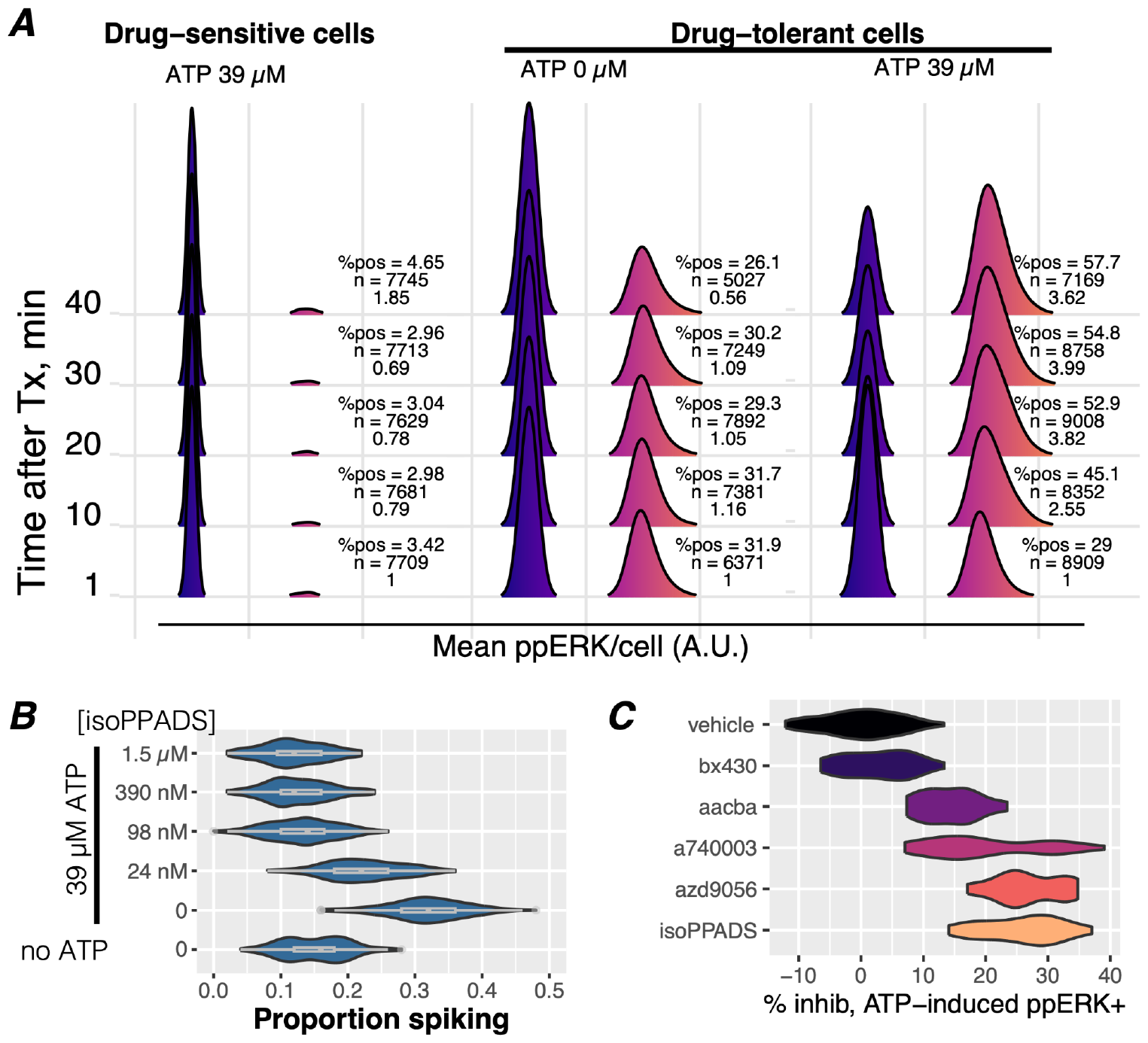
Exogenous ATP enhances Ca^2+^ spiking and ppERK activation via P2X7 receptors in drug-tolerant cells. *A)* Immunofluorescence detection of dual-phosphorylated ERK (ppERK) was performed on drug-sensitive (60 min BRAFi) and drug-tolerant (3 day BRAFi) cells treated with 39 μM of exogenous ATP or vehicle control (0 μM). Cells without detectable ppERK staining were assigned values of –0.5 and are represented by the blue distributions. Intensity of ppERK staining is represented by the fuschia-colored distributions. A ppERK intensity score is calculated from the percent positive cells and the intensity distributions of ppERK-positive cells and normalized to the 1 min time point. The ppERK intensity score, the number of cells in each condition, and the percentage of ppERK positive cells (%pos) are indicated to the right of each pair of distributions per condition. The ppERK intensity score and %pos increased in a time-dependent manner in drug-tolerant but not drug-sensitive cells. *B)* [Ca^2+^]_cyt_ spiking activity was assessed in Calbryte-loaded A375 cells treated with BRAFi for 3 days without or with 39 μM ATP stimulation and without or with pretreatment of the P2Xi isoPPADS at the indicated concentrations. The addition of ATP significantly increased the proportion of spiking cells, which was inhibited by the addition of isoPPADS. *C)* A panel of P2X inhibitors (100 nM each) or vehicle were preincubated on drug-tolerant A375 cells, treated with 39 μM ATP for 30 min, and stained for ppERK activity. Inhibition of ATP-induced ppERK staining intensity is shown as a percent of inhibition relative to the vehicle control and shows that all P2X inhibitors that affect P2X7 activity inhibit ATP-induced ppERK, in contrast to the P2X4 inhibitor (bx430), which had no effect.

Our studies have thus far indicated that adaptation to targeted BRAFi treatment alters multiple axes of Ca^2+^ signaling, at least one of which is relevant to the drug-tolerant state via ppERK re-activation (Fig. 2D, 2E). Moreover, this effect is not limited specifically to BRAFi since the [Ca^2+^]_cyt_ spikes occur despite co-treatment with a MEK inhibitor (BRAFi + MEKi) (Supplementary Fig. S1). Altogether, these findings demonstrate that drug-tolerant melanoma cells overexpress purinergic receptor P2X7 (Fig. 2A, 2B) that are functionally active in inducing [Ca^2+^]_cyt_ spikes (Fig. 1B, 2C, 3B) and are responsive to ATP and that the inhibition of P2X7-mediated [Ca^2+^]_cyt_ spikes reduces ppERK reactivation in drug-tolerant cells (Fig. 3C).

### Ca^2+^-mediated activation of ERK is potentiated in drug-tolerant cells

Previous work has demonstrated an increased ability of receptor tyrosine kinases to potentiate ERK activation in the presence of BRAFi due to the release of negative feedback^42^. This is an accepted mechanism of BRAFi bypass that underscores the adaptability of melanoma cells to treatment, presumably in a non-genetic fashion. Similarly, Ca^2+^ signaling in BRAFi tolerant cells may be a crucial and parallel component of this mechanism, reflecting phenotypic adaptation that leads to efficient Ca^2+^-mediated reactivation of ERK. To directly test this possibility, we compared short- (60 minutes) versus long-term (3 days) BRAFi treatment, followed by a time-course exposure to the Ca^2+^ ionophore, ionomycin. Ionomycin drives e[Ca^2+^] into the cytoplasm, thereby inducing a Ca^2+^ signal in all cells. Following treatment, cells were fixed and immunostained for ppERK to quantify Ca^2+^-mediated activation of ERK (Fig. 4A). We found that in drug-tolerant cells, ERK phosphorylation was stimulated at lower concentrations of ionomycin, reached much higher levels at a faster rate, and stayed elevated longer than drug-sensitive cells (Fig. 4B). Furthermore, the maximum percentage of cells staining for ppERK in the long-term treated population (>40%) was higher than the short-term treated population (<25%) (Fig. 4B), suggesting that more of the population is capable of Ca^2+^-mediated activation of ERK. The addition of another Ca^2+^ mobilizing agent, cyclopiazonic acid (CPA), which causes the passive release of ER Ca^2+^, also leads to reactivation of ppERK (Supplementary Fig. S12) in drug-tolerant but not drug-sensitive cells. Consistent with our observation of reduced ER Ca^2+^ stores, CPA activates ppERK to a lesser extent than ionomycin, which is likely a reflection of the amount of Ca^2+^ flux that occurs. These results support the conclusion that drug-tolerant cells, adapted over several days, have an enhanced ability to activate the MAPK pathway via Ca^2+^ signaling.

**Figure 4.**
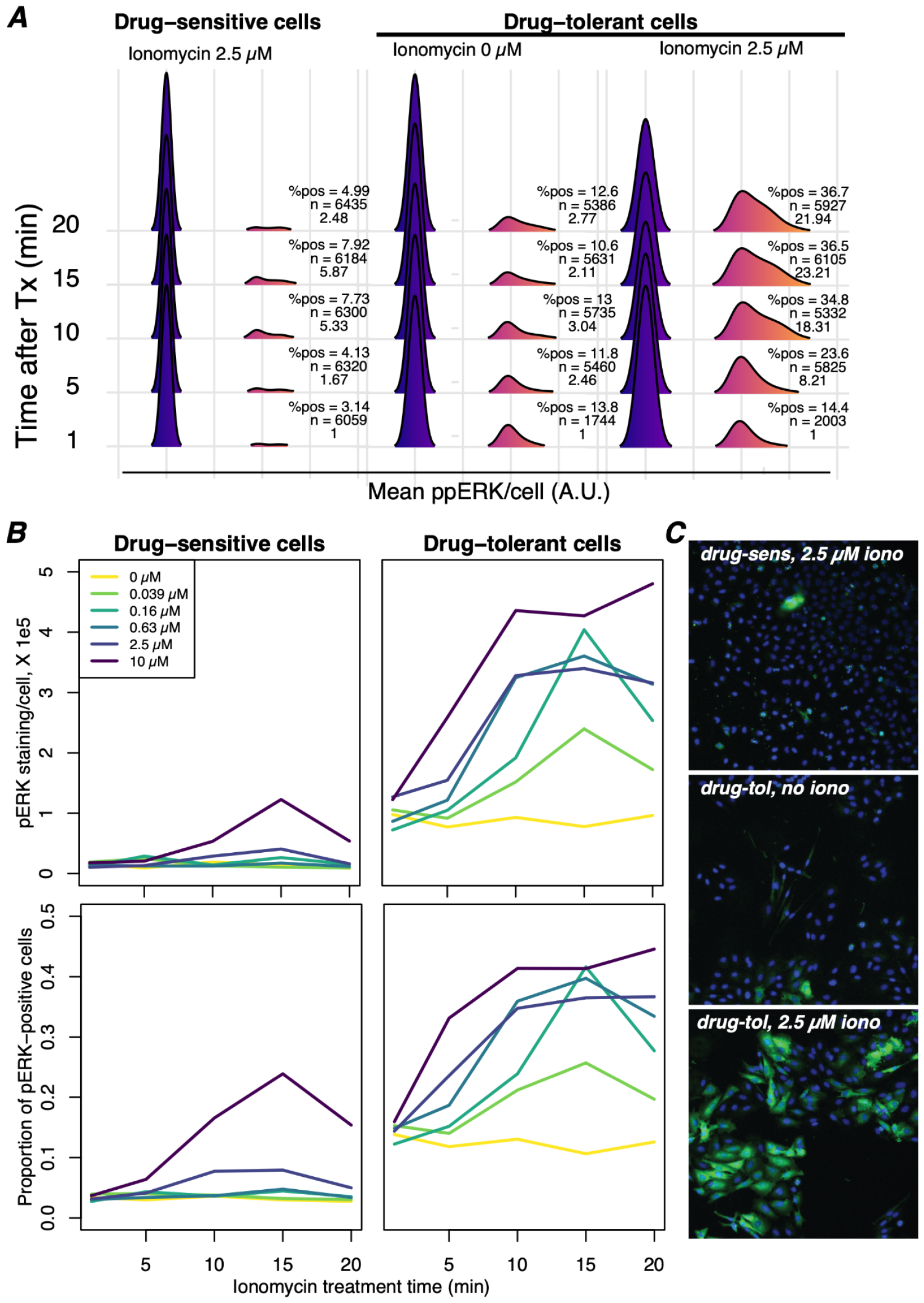
Ca^2+^ mobilization induces greater ppERK activation in drug-tolerant than drug-sensitive cells. *A)* Immunofluorescence detection of ppERK was performed on drug-sensitive (60 min BRAFi) and drug-tolerant (3 day BRAFi) cells in the presence of the Ca^2+^ ionophore ionomycin (iono). The percentage of positive cells and mean ppERK intensity per cell increased in a time-dependent manner after the addition of ionomycin in drug-tolerant cells but not in drug-sensitive cells. *B)* Mean ppERK staining intensity per cell *(top)* and the proportion of cells staining positive for ppERK *(bottom)* across a time series and concentration range of iono. Iono induced ppERK staining in drug-sensitive cells with concentrations at or above 2.5 μM, whereas ppERK was activated with as little as 39 nM iono in drug-tolerant cells, demonstrating a particular sensitivity to Ca^2+^ mediated activation of ppERK. ppERK staining increases at the earliest time point (5 minutes) in drug-tolerant cells, but not drug-sensitive cells, indicating that drug-tolerant cells respond more rapidly to Ca^2+^. Mean staining intensity in positive cells were significantly elevated at all tested concentrations of ionomycin in drug-tolerant cells but only the highest concentration in drug-sensitive cells. Moreover, iono activated ppERK in a greater proportion of drug-tolerant cells and did not diminish during the time series at higher concentrations of iono, in contrast to drug-sensitive cells. *C)* Representative images of ppERK staining (green) with Hoechst-stained nuclei (blue) under the indicated conditions, each after 15 min of treatment. All images were captured with the identical settings and have been set to the same intensity range for visualization.

## DISCUSSION

Here we identify a previously unrecognized signaling network responsible for ERK reactivation in drug-tolerant cells based upon the following findings. 1) BRAFi induces robust cell-intrinsic [Ca^2+^]_cyt_ spikes in drug-tolerant cells, which become more pronounced with treatment time and correspond to increased ppERK levels. 2) Increases in P2RX7 transcript abundance is consistent across multiple cell lines and with increased Ca^2+^ flux. 3) [Ca^2+^]_cyt_ spikes are dependent on e[Ca2+], and can be blocked by P2X7 inhibitors. 4) Inhibition of [Ca^2+^]_cyt_ spikes reduces ERK reactivation in drug-tolerant cells. 5) Drug-tolerant cells activate MAPK with much greater sensitivity to Ca^2+^ flux.

An intriguing feature of this network is that ATP, which is virtually ubiquitous in living systems, is the ligand that can initiate [Ca^2+^]_cyt_ spikes via P2X receptors. ATP is abundant in the tumor microenvironment and is released by dying cells, thereby implicating the death of drug-sensitive cells as a source of trophic stimuli that leads to ERK reactivation and drug-tolerant behavior. In direct support, we found that cell lysates stimulated an increase in ppERK, similar to ATP, ionomycin, and CPA (Supplementary Fig. S13). Such a mechanism immediately offers an explanation of the inherent efficacy issues of BRAFi treatment in BRAF-mutant melanoma. That is, both *in vitro* and *in vivo* BRAFi-induced cell death may simultaneously favor survival and proliferation of adapted drug-tolerant cells by the release of ATP, a mitogenic ligand. Accordingly, we found the percentage of cells with [Ca^2+^]_cyt_ spiking and those capable of Ca^2+^ mediated activation of MAPK increased with treatment time in drug-tolerant populations (Fig. 1C and 4C), suggesting a selective advantage is conferred by mitogenic [Ca^2+^]_cyt_ spikes (Fig. 2D, 3A, and 4A).

Parallel to ATP release from dead and dying cells, Pannexin-2, also enriched in drug-tolerant cells (>2 fold) (Fig. 2A), may form a cell-to-cell positive feedback loop amongst adjacent drug-tolerant cells, since it reinforces purinergic receptor activation by ATP release^65^. It is tempting to speculate ATP may be the diffusible ligand responsible for ppERK pulses in drug-tolerant melanoma cell clusters, as described by Gerosa et. al.^66^

Previous reports have linked melanoma tolerance to BRAFi with RTK activity (e.g., AXL, FGFR, NGFR, EGFR^9,67^) and ERK reactivation^66,68^. However, no clinically effective RTK combination in BRAF-mutant melanoma has been identified. In contrast, targeting EGFR in BRAF-mutant colorectal cancer^69,70^ has been successful, underscoring the significance and possibility of a variety of context-dependent mechanisms of MAPK reactivation. These RTK-based mechanisms of ERK reactivation are ligand dependent, raising the question of possible exogenous sources for these ligands *in vitro* and *in vivo*. In contrast, in the ppERK reactivation pathway we describe here, the ATP ligand may be released by cells dying from BRAFi, which can then be used for survival by neighboring cells to reactivate ERK via P2X receptor [Ca^2+^]_cyt_ spikes. It is possible that P2X7 signaling activity could also underlie RTK signaling activity that may occur in the drug-tolerant state via transactivation mechanisms^52,60,71–79^.

It has been convincingly shown that drug tolerance is a heterogeneous collection of distinct cell states^15,25–27,80^. Our observation of diverse patterns of spiking (Fig. 1B) in drug-tolerant cells suggests a possible correlation with distinct biochemical events or cell states within tolerant cell populations^3^. Additional work to establish such connections may distinguish amongst separate mechanisms of tolerance resulting from specific [Ca^2+^]_cyt_ spike patterns. A limitation of our study is the exclusive focus on ERK reactivation downstream of purinergic [Ca^2+^]_cyt_ spikes. There are a plethora of other cellular functions in which Ca^2+^ plays a role, independent of ppERK reactivation, that may be relevant to drug-tolerance, including bioenergetics, metabolism, stress response, and cell cycle progression^39^. Furthermore, activation of Ca^2+^ signaling beyond purinergic cation channels should also be considered since P2X7 inhibitors did not inhibit all [Ca^2+^]_cyt_ spikes or ppERK staining, indicating that other Ca^2+^ channels may also contribute to drug tolerance (Fig. 1A, 2A, and Supplementary Fig. S6). However, it is possible P2X7 inhibitors do not block all P2X7 channel activity but may instead modulate Ca^2+^ channel properties (i.e., result in altered spike patterns) without completely ablating spikes. To address these possibilities, more sophisticated methods to analyze [Ca^2+^]_cyt_ spiking patterns are needed to reveal more nuanced relationships. For example, isoPPADS appears to have a greater inhibition of ppERK (∼25%) at lower concentrations (Fig. 2E) then would be expected from the complete ablation of [Ca^2+^]_cyt_ spikes observed in Fig. 2C. Intriguingly, the same cannot be said for AZD9056, demonstrating the context dependence of these inhibitors. It will be relevant to elucidate the relationships between the other P2X7 inhibitors, [Ca^2+^]_cyt_ spike properties, and ppERK activity with additional experiments and improved analysis methods of [Ca^2+^]_cyt_ spikes, which has been enigmatic in the field of Ca^2+^ signaling. Nonetheless, given the enhanced ability of Ca^2+^ to activate ERK in the drug-tolerant state, it will be worthwhile to elucidate if these [Ca^2+^]_cyt_ signals, regardless of the source, underlie most of ERK reactivation and, therefore, drug-tolerant behavior. Accordingly, we have observed that drug-tolerant cells do not tolerate the absence of extracellular Ca^2+^ (Supplementary Fig. S14), indicating that inhibiting all [Ca^2+^]_cyt_ spikes may be a viable therapeutic approach.

Raj and co-workers described a jackpot effect^9,68^, whereby a stochastic combination of non-mutational (i.e., epigenetic) events generate rare cells in BRAF-mutant melanoma cultures under BRAFi. These jackpot cells are implicated in ERK reactivation-dependent escape from BRAFi and appear within a time frame similar to the drug-tolerant idling state we describe. However, by definition the jackpot cells are a rare occurrence. In contrast, the drug tolerance we describe here is a cell population phenomenon. That is, a large portion of the population adapts to the BRAFi stress (Supplementary Movie M1), and no selection of rare clones takes place^32^. Thus, idling and jackpot descriptions are distinct, although possibly related. It would be interesting to determine whether [Ca^2+^]_cyt_ spikes appear at any stage in jackpot cells and/or play a causative role in their development. Spencer and co-workers reported that drug-tolerant cells dividing in the presence of BRAFi tend to have increased ER stress response signaling in comparison to non-dividing cells. ER stress-driven autophagy has previously been shown to be a survival mechanism utilized by melanoma cells under therapeutic treatment^81,82^. Whether these findings are related to the [Ca^2+^]_cyt_ spikes in idling cells is not known. However, ER stress signaling can be induced by insufficient ER Ca^2+^ stores, and our data indicate that drug-tolerant cells have diminished ER Ca^2+^ content.

In normal skin, UV light-induced ATP release from keratinocytes activates P2X7 receptors on melanocytes to induce melanogenesis and melanosome transfer to keratinocytes^83^. It remains to be determined whether the increased expression and activity of P2X7 in melanoma under BRAFi may be related to this physiological function. Along these lines, BRAFi is known to induce a range of melanocytic differentiation states^3^. Moreover, this possible relationship may be reflected in the relatively high but wide range of P2RX7 transcript abundance in melanoma compared to other tumor types (Supplementary Fig. S4).

Purinergic signaling is an expansive area of active research. It is especially studied in the nervous system, where it can participate in synapse formation, action potentials, and various neuropathologies. In the immune system, purinergic receptors and associated surface molecules modulate T cell activation and are key players in inflammation^36^. In comparison, purinergic signaling has been less studied in cancer, although it has gained attention in recent years. Immunosuppressive effects and modulation of host–tumor cell interactions mediated via purinergic receptors and ATP in the tumor microenvironment was reviewed^36^. Here, we put forward the novel perspective that ATP-driven activation of P2X receptors may be an adaptive mechanism of escape from targeted therapy (e.g., BRAFi), by promoting the rise of drug-tolerant cells.

In response to ATP, drug-tolerant cells maintain a sustained concentration of [Ca^2+^]_cyt_ compared to drug-naive cells (Supplementary Fig. S11). This observation may be worthy of attention, and could be due to several non mutually exclusive possibilities. One is differential channel gating (i.e., increased open probability) mediated by post-transcriptional (e.g., alternative splicing) or post-translational mechanisms^84^. Alternatively, Ca^2+^ clearance may be reduced, which is consistent with our findings (Supplementary Fig. S15). Regardless, tolerant cells may be intrinsically more prone to maintain higher levels of [Ca^2+^]_cyt_, thus enhancing Ca^2+^ signals that promote adaptive response. Clarifying this may provide actionable implications for inhibiting tolerance.

Recently, P2X7 activation was associated with improved therapy responses in NRAS-mutant melanoma^85^, presumably by reducing the rise of resistant cells. This is contrary to the role of purinergic signaling in BRAF-mutant melanoma that enhances drug tolerance via ppERK reactivation during BRAFi therapy. It is nonetheless possible that these discordant findings point to fundamentally distinct biology of mutations occurring at different sites in the MAPK pathway. More work is needed to reconcile these apparently conflicting observations. For instance, it is worth determining whether in one case there is much higher P2X7 activation than the other, such that the level of P2X7 activation and thus Ca^2+^ flux could cause opposite outcomes. Indeed, our findings indicated that excessive ATP stimulation in drug-tolerant cells reduces viability (not shown), which is in agreement with prior studies showing prolonged exposure of cells to high concentrations of ATP induce cell death^86^.

Although our results implicate P2X7 as a primary driver of the increased Ca^2+^ signaling in drug tolerance, other Ca^2+^ regulatory genes are likely to contribute to the cellular responses. For example, P2X-induced [Ca^2+^]_cyt_ spikes may initiate Ca^2+^-induced Ca^2+^ release through ryanodine receptors on the ER, which are facilitated by low-voltage-gated calcium channels (CACNA1C and CACNA1D) that are known to activate RYR in heart muscle and were observed to increase in expression in drug-tolerant cells (Fig. 2A). The release of ER Ca^2+^ in response to [Ca^2+^]_cyt_ spikes could explain the reduced ER Ca^2+^ content and be a contextual difference that explains the apparently discordant findings in NRAS and BRAF mutant melanoma.

The MAPK signaling architecture of BRAFi tolerant melanoma cells is uniquely adapted to efficiently use Ca^2+^ signaling. This property suggests that targeting Ca^2+^ signaling may be key to increasing durability of treatment while maintaining enough target specificity for treatment tolerability. Such approaches are desirable because they could reduce patient suffering in comparison to the standard-of-care combinations with MEK inhibitors, which come with significant side effects. Indeed, clinical trials demonstrate P2X receptor inhibitors are well tolerated in patients, providing confidence that targeting P2X signaling in combination with BRAFi could be a promising strategy to combat drug tolerance.

## METHODS

### Cell culture

BRAF mutant melanoma cell lines were grown in DMEM/F12 (Gibco Ref 11330-032) supplemented with 10% fetal bovine serum and 1% pen/strep, at 37°C and 5% CO_2_ as described previously {Paudel; Hardeman}.

### Reagents

The intracellular calcium dyes, Calbryte-520-AM (AAT Bioquest, 20650) and Fura-2-AM (AAT Bioquest, 21023), were individually dissolved in DMSO and frozen in single use aliquots to prevent freeze thaw. The drugs in Supplementary Table 1 were dissolved in indicated solvent (DMSO or water), aliquoted to single use tubes, and stored at -20°C. Ionomycin calcium salt (Cayman Chemical, Ref 11932) or cyclopiazonic acid (CPA) (Cayman chemical, Ref 11326) were dissolved in DMSO to 10 mM and 50 mM respectively, aliquoted into single use tubes, and stored at -20°C for no longer than 3 months.

### Drug treatment

Melanoma cells were detached from flasks using TrypLE (Thermo Fisher), counted, diluted, and seeded into 384-well plates (Greiner 781091) or 35 mm imaging dishes (Cellvis D35-20-1.5P), in 30 μL or 2 mL volumes, respectively. The number of cells seeded is chosen for each condition to achieve approximately 60–70% confluency at the time of experimental observation. For 384-well plates, drug- or vehicle-containing medium was added at 2X concentration to existing media the following day (approximately 18–24 hours later). For 35 mm dishes, existing media was aspirated and 2 mL of drug- or vehicle-containing medium was added the following day (approximately 18–24 hours later). In experiments with long term treatment, medium was aspirated and fresh vehicle- or drug-containing medium was added every 3–4 days. The research analog of vemurafenib, PLX4720, was used for BRAF inhibition, except where noted.

### Dye loading for imaging of cytoplasmic calcium

#### Fura-2-AM

For experiments using Fura-2, unless otherwise noted, Imaging Buffer was Ca^2+^ containing Hanks Balanced Salt Solution (Ca^2+^:HBSS) (Gibco 14025-092) containing any relevant drugs (i.e. PLX4720). To cells in 35 mm dishes, 1 mL of medium was removed and Fura-2-AM was added to 4 μM (2X) concentration before adding back to the plate to achieve a 2 μM final concentration. Plates were incubated at 37°C, 5% CO_2_ for 30 minutes. After dye loading, the media was aspirated and the plates were washed 3X with 2 mL of Imaging Buffer warmed to 37°C. After the final wash, 2 mL of Imaging Buffer was added to the dish.

#### Calbryte-520-AM

For experiments using Calbryte, unless otherwise noted, Imaging Buffer was phenol red- and serum-free culture medium (DMEM/F12, Gibco Ref 11039-021) containing any relevant drugs (i.e. PLX4720). Cells plated to 384-well plates were loaded with 2 μM Calbryte-520-AM diluted in culture media containing relevant drugs (i.e. PLX4720). To begin dye loading, media from cell plates were aspirated down to 20 μL with a BioTek ELx405 plate washer (ELX plate washer), followed by addition of 20 μL 2X dye-containing medium (warmed to 37°C) for a final concentration of 2 μM Calbryte-520-AM. The plates were incubated at 37°C, 5% CO_2_ for 25 minutes before 20 μL of medium was aspirated and wells were washed 3X with 80 μL of Imaging Buffer. Before imaging, 20 μL of Imaging Buffer was added to achieve a final volume of 40 μL. When Ca^2+^ containing and Ca^2+^ free imaging conditions were compared (as in Fig. 1D), the imaging buffers were Ca^2+^:HBSS (Gibco 14025-092) and Ca^2+^ free HBSS (Gibco 14175-095), respectively.

### Live-cell imaging of spontaneous cytoplasmic calcium spikes

#### Fura-2-AM

After dye loading/washing, imaging dishes were moved to a heated stage. Fura-2-AM fluorescence (Ratio 340Ex/380Ex-535Em; 340/380 ratio) was measured every 5 seconds as an indicator of intracellular Ca^2+^; when the intracellular concentration of Ca^2+^ increases, so does the 340/380 ratio. Imaging was performed for a period of 40 minutes with a Nikon Eclipse Ti2 microscope using a 10X objective and equipped with a Photometrics Prime 95B 25mm sCMOS Camera. For Ca^2+^ free conditions, cells were washed and imaged in Ca^2+^ free HBSS (Gibco 14175-095).

#### Calbryte-520-AM

After dye loading/washing, plates were moved to an ImageXpress Confocal HT.ai imager with environmental controls (37°C, 5% CO2, humidity) for 15 minutes to allow for equilibration. When purinergic inhibitors were included in the imaging conditions, they were added at 2x concentration immediately after the final wash step and before the 15 minute equilibration. If additional inhibitors were not included, then 20 μL of Imaging Buffer were added to wells, as described above. Calbryte-520 signal was imaged simultaneously across 8 wells at an interval of 5 seconds within each well to detect spikes in cytoplasmic calcium levels. Imaging was performed with a FITC filter set, using a 470 nm laser light source (89North). Imaging was performed for 40 minutes, unless otherwise noted. To link Calbryte images to ppERK data within individual cells, at the conclusion of Calbryte imaging, cells were fixed with 4% paraformaldehyde (PFA) (Thermo Scientific Chemicals, 043368.9M) at room temperature for 20 minutes followed by 3X washes with 80 μL PBS. ppERK staining was performed as described below.

### Live-cell imaging of ATP-stimulated cytoplasmic calcium spikes

#### Calbryte-520-AM

Cells were loaded with Calbryte-520-AM as described above and ATP-containing Imaging Buffer (maintained at 37°C) was added to each well immediately before imaging on an ImageXpress Micro XLS automated microscope (Molecular Devices) using a 10X objective and a Lumencor Sola solid state light source. Calbryte-520 fluorescence was excited and detected using a Semrock GFP filter set (GFP-3035B) consisting of a 472/30 excitation filter, 442-488 / 502-730 dichroic, and 520/35 emission filter. One site per well was captured using a 4.2MP widefield scientific cMOS camera with a camera binning of 1. One well at a time was imaged at 5 second intervals for 5 minutes. For experiments using the P2X inhibitor, isoPPADS, the drug (or vehicle control) was serially diluted in the Imaging Buffer and preincubated on cells for 15 minutes at 37°C, 5% CO_2_ after the final wash step.

### Live-cell imaging of Store Operated Calcium Entry (SOCE)

#### Fura-2-AM

SOCE assays were conducted, as described previously^87^, in three phases consisting of 1) an equilibration period in Ca^2+^-free HBSS, 2) release of Ca^2+^ from the endoplasmic reticulum (ER), and 3) the induction of SOCE activity. Changes in cytoplasmic Ca^2+^ levels were measured with Fura-2-AM, as described above. Briefly, cells were dye loaded, washed, and incubated in Ca^2+^ free HBSS before imaging. Phase 1) Ca^2+^-free HBSS was perfused over cells for 10 minutes of imaging to allow equilibration and to acquire a baseline. Phase 2) Ca^2+^-free HBSS buffer containing 50 μM of the sarco-endoplasmic Ca^2+^ ATPase (SERCA) inhibitor, cyclopiazonic acid (CPA), was perfused over cells for 8 minutes. During this phase, CPA treatment leads to the release of free Ca^2+^ within the endoplasmic reticulum (ER), allowing for the quantification of ER Ca^2+^ stores and subsequent cytoplasmic Ca^2+^ clearance. Release of ER Ca^2+^ stores results in activation of SOCE channels, however, they do not contribute to the Ca^2+^ signal until extracellular Ca^2+^ is added to the assay in the third phase. Phase 3) Perfusion with Ca^2+^:HBSS (+50 μM CPA) allows influx of Ca^2+^ through activated SOCE channels, and subsequent quantification of SOCE activity on a cell by cell basis. This final component of the assay proceeds for an additional eight minutes before imaging is stopped. For all phases, cells were perfused at a flow rate of 2 mL/min. Quantifications of ER Ca^2+^ content and SOCE activity are calculated by taking the integral of each cell trace within the respective phases, followed by baseline subtraction.

### ppERK staining

Cells were fixed with a final concentration of 4% PFA at room temperature for between 20 and 40 minutes. Cells were washed 3X with 80 μL PBS using an ELX plate washer, leaving 15 μL after the final wash and aspiration step. 15 μL of 2X primary antibody were added (Phospho-p44/42 MAPK (Erk1/2) (Thr202/Tyr204) (D13.14.4E) XP Rabbit mAb #4370; Cell Signaling, Ref 4370S), generating a final concentration of 1:800 (3% goat serum, 0.3% Triton X100). Plates were incubated at 4°C overnight. The next morning cells were washed 3X with PBS and 15 μL of 2X secondary antibody (Donkey anti-Rabbit IgG (H+L) Highly Cross-Adsorbed Secondary Antibody, Alexa Fluor™ 488; Invitrogen, Ref A21206) were added to the remaining 15 μL of PBS to generate a final concentration of 1:1000 (3% goat serum, 0.3% Triton X100). Plates were incubated for 2 hours protected from light at room temperature before washing 3X with PBS, and 15 μL of 2X Hoescht 33342 (Thermo Scientific, 62249) (1.7 μM final concentration) were incubated on cells for 5 minutes before washing 3X with PBS. Plates were sealed with an aluminum plate sealer. Imaging of nuclei and ppERK signal was performed using an ImageXpress Micro XLS automated microscope (Molecular Devices) with a 10X objective and a Lumencor Sola solid state light source. Alexa-488 fluorescence was excited and detected using a Semrock GFP filter set (GFP-3035B) consisting of a 472/30 excitation filter, 442-488 / 502-730 dichroic, and 520/35 emission filter. Four sites per well were captured using a 4.2MP widefield scientific cMOS camera with a camera binning of 1. Multiple plates were loaded to a plate carousel (ThermoScientific), shuttled to/from the ImageXpress via a ThermoScientific CRS F3 robot arm, and automatically run through the Momentum scheduling software (Thermo). Data was quantified using MetaXpress multiwavelength cell scoring. Primary alone or secondary alone antibody controls were captured to quantify background signal.

### P2Xi effects on ppERK

P2X inhibitors were diluted to 3X concentrations in warmed, serum-free cell culture media and serially diluted within 384 well reservoir plates (Greiner, Ref 781280) using a Bravo automated pipette liquid handler (Velocity 11/Agilent). Media from cell plates were aspirated using an ELX plate washer, leaving 20 μL volumes. A Bravo liquid handler added 10 μL of media containing each 3X concentration of drug (or vehicle) to respective wells, and plates were returned to the incubator for an hour before 10 μL of 16% PFA (4% final concentration) were added to each well using a ThermoScientific Multi-Drop Combi liquid dispenser to fix cells. Cells were incubated for 20 minutes at room temperature before washing 3X with PBS. Staining for ppERK was performed as described above. All conditions were performed in technical triplicate on the same plate.

### Ca^2+^ mobilization effects on ppERK

Ionomycin calcium salt (Cayman Chemical, Ref 11932) or CPA (Cayman chemical, Ref 11326) stocks were diluted to 2X the max desired concentration in serum free culture media, then serially diluted within 384 well reservoir plates using a bravo automated pipette to generate a drug addition plate. Media was aspirated from cell plates using an ELX plate washer immediately before stimulation, leaving 15 μL/well. 15 μL from the drug addition plate were stamped onto the cell plate and incubated at room temperature for the duration of the time series. At each indicated time point, 10 μL of 16% PFA (to 4% final concentration) were added to the corresponding wells. At the last time point, PFA was added and incubated an additional 20 minutes at room temperature before the entire plate was washed with 3X with PBS using an ELX plate washer. Staining for ppERK was performed as described above. All conditions were performed in technical triplicate on the same plate.

### ATP solution

ATP (MedChemExpress, Ref HY-B2176) was dissolved in a solubilization buffer consisting of 200 mM HEPES and 3 mM CaCl_2_ in DEPC H_2_O, to create a stock solution with a concentration of 300 mM ATP. This solution was buffered to a pH of 7 using 1 M KOH and 5 M NaOH; final concentrations were approximately 6 mM KOH and 600 mM NaOH. This stock was frozen in aliquots and stored at -20°C for no more than 3 months. Aliquots were diluted in the relevant experimental buffer to generate the indicated final concentrations.

### ATP stimulation of ppERK

The ATP solution described above was diluted to a max concentration in serum free culture media and serially diluted using the Bravo instrument, as described above. Media were aspirated to 15 μL/well using an ELX plate washer immediately before stimulation. For each time point, 15 μL of the 2X ATP-containing solution were added to the cell plate and cells were maintained at 37°C and 5% CO_2_ to achieve 1, 10, 20, 30 or 40 min ATP treatment time (reverse time series where all treatment times are completed simultaneously). After incubation, 10 μL of 16% PFA (4% final concentration) were added to all wells using a ThermoScientific Multi-Drop Combi and incubated at room temperature for 20 minutes. Cells were washed 3X with PBS, before primary staining for ppERK as described above. All conditions were performed in technical triplicate on the same plate.

For experiments with P2X inhibitors, titrations of each P2X inhibitor (or vehicle alone) were generated by serial dilution with the Bravo in serum-free culture media. Media were aspirated, leaving 20 μL/well, immediately prior to the addition of 5 μL of each P2Xi solution (6X) followed by incubation at 37°C, 5% CO_2_ for 30 minutes. After the P2Xi preincubation, 5 μL of 6X ATP (or vehicle) were added to each well to generate a final concentration of 39 μM ATP. The plates were returned to 37°C, 5% CO_2_ for 30 additional minutes. Cells were then fixed and stained for ppERK as described above.

### Bulk RNAseq

#### Drug treatment

Four different BRAF-mutant melanoma cell lines (A375, SKMEL28, SKMEL5, and WM88) were seeded overnight into 10 cm dishes in their standard culture medium and treatment with 8 μM PLX4720 was initiated the following day on three separate occasions (biological triplicate). Media containing PLX4720 were replaced every 3 days up to 8 days of treatment. Media were aspirated and cells were kept on ice while washing with 25 mL of ice-cold PBS followed by aspiration and addition of 1 mL of ice-cold PBS into which cells were gently scraped into suspension. PBS cell suspension was pipetted to a pre-chilled to 1.5 mL RNase-free snap lid tube (Zymo Research, Cat C2001-100; used for all downstream sample handling). Cells were pelleted by centrifugation at 500x*g* @ 4°C for 5 minutes and washed again with 1 mL ice-cold PBS. The remaining cells in the tube were immediately snap-frozen in liquid nitrogen and stored at -80°C until all samples were collected. Drug naive cells were collected similarly in biological triplicate, except cells were seeded and collected two days later such that drug-treated and drug-naive conditions had approximately the same number of cells per plate upon harvesting.

#### RNA isolation

The samples were processed in groups of eight with one biological replicate from each cell line processed at a time (24 samples total). Samples were briefly thawed on ice, resuspended in 1 mL Trizol, and vortexed. Samples were incubated at room temperature for 5 minutes before 200 μL of chloroform was added. Samples were vigorously shaken for 10 seconds then moved to ice for 3 minutes. Samples were centrifuged at 4°C and 13,000x g for 15 minutes. Approximately 400 μL of the top aqueous phase was moved to 500 μL of cold isopropanol in a new 1.5 mL tube and incubated at -20C for 1 hour. Samples were centrifuged at 4°C and 13,000x*g* for 15 minutes to pellet RNA. Isopropanol was removed and 1 mL ethanol was added to wash the pellets. Samples were again pelleted at 4°C and 13,000x*g* for 15 minutes before ethanol was removed and samples allowed to air dry. Samples were resuspended in DEPC-treated H_2_O and treated with Qiagen RNase free DNAse (ref #79254), per the protocol’s recommendation (15 minute room temperature incubation). After DNAse treatment, RNA was reisolated using the same procedure. 0.5 mL of Trizol and 100 μL of chloroform were added to the samples and shaken vigorously. Samples were placed on ice for 3 minutes before being centrifuged at 4°C and 13,000x g for 15 minutes. Approximately 250 μL of aqueous phase was transferred to 300 μL of cold isopropanol and incubated at -20C for one hour. RNA was pelleted at 4°C and 13,000x g for 15 minutes, isopropanol decanted, and 1 mL of ethanol added to wash the samples before being centrifuged again at 4°C and 13,000x g for 15 minutes. Ethanol was decanted and pellets air dried before being resuspended in 30 μL DEPC-treated H_2_O. RNA concentration was measured by nanodrop and aliquots sent off for RIN analysis of RNA quality.

#### Transcriptomic Sequencing

Illumina NovaSeq 6000, stranded mRNA (poly-A), paired ends 150 bp reads.

#### Sequence processing

Read quality control was performed using FastQC v0.11.9 (https://www.bioinformatics.babraham.ac.uk/projects/fastqc/). Reads were aligned to the human reference genome (GRCh38.p14) using STAR version 2.7.7a^88^ and GENCODE version 21^89^ using the default settings and transcript abundance was determined using STAR’s “quantMode.” Alignment quality control was performed with Picard

{http://broadinstitute.github.io/picard/}. Links to scripts can be found in Supplementary Information.

### Image processing

#### ppERK images

Molecular Devices MetaXpress Multiwavelength Cell Scoring (https://www.moleculardevices.com/sites/default/files/en/assets/data-sheets/dd/img/metaxpress-software-multi-wavelength-cell-scoring-application-module.pdf) was used to quantify object-level information from the images, which were then used to determine the number of cells staining positive for ppERK and signal intensities of positive cells. Nuclei were identified with the Hoescht nuclear channel and cytoplasmic cell boundaries were identified from the ppERK antibody fluorescence channel. Cell masks were generated by associating nuclei and ppERK signal intensities greater than background fluorescence (i.e. absence of primary antibody). Cells without intensity values above the background threshold were characterized as negative for ppERK staining.

#### Fura-2 images

Cell masks were generated using Nikon’s NIS analysis platform from images obtained using 340 nm excitation. Mean signal intensity within the cell masks for each excitation wavelength were quantified at each time point across the time series. Ratios of signal intensities from 340 nm and 380 nm excitation (340/380) were used as a measure of the proportion of Ca^2+^-bound to Ca^2+^-free dye, providing a direct relationship to intracellular Ca^2+^ levels.

*Calbryte images:* Images obtained from an ImageXpress Micro XLS (16-bit grayscale images of 1080 X 1080 pixels) and annotation files describing the time and well position of each image were assembled using custom R code (available at https://github.com/darrentyson/IXprocess). Time series image stacks were assembled for each well of data and exported as a multipage TIF file for further processing. The first image of each well’s time series was used to obtain a labeled mask image with adjacent/contiguous pixels defining each object being assigned a unique value (region ID) using DeepCell pretrained model for segmentation (www.deepcell.org/predict10.1038/s41592-020-01023-0). The labeled segmentation masks were then used to quantify integrated intensity values for each object from each image of the relevant calbryte time series using custom Python scripts (all code for performing these steps is available on GitHub).

### Data analysis

All code used for analyses are provided in a public git repository available on GitHub (https://github.com/QuLab-VU/Stauffer_et_al_2024).

*Differential expression of transcript abundance* was performed on transcript counts from four melanoma cell lines in R (v4.3.2) using DESeq2^90^ (v1.42.0). Data from A375 cells was used as reference to facilitate comparison to results in Gerosa et al., 2020. Enrichment of genes associated with drug tolerance compared to the drug-naive state were identified using the “contrast” selection of the DESeq2 results object. Gene ontology term enrichment was performed using “clusterProfiler”^91^ on differentially expressed genes filtered with Benjamini-Hochberg-adjusted p-value of 0.05 and 1.5-fold change of expression as cutoffs.

#### ppERK

Only objects of total area between 50 and 4500 pixels^2^ with integrated intensity values of < 5e6 were considered. Proportions of positive cells were generated by dividing the number of positive cells by the number of total cells per condition and were reported as percentages or used to calculate percent inhibition versus control by setting the proportion of positive cells in the control condition as the denominator. We conservatively estimated the dose–response relationships of P2Xi on ppERK staining by assuming drug concentrations of 1 nM for each drug had no effect (i.e., were similar to vehicle control) to allow log-logistic model regression.

To estimate the effects of P2Xi on ATP-induced ppERK, percentage inhibition was calculated as the percent reduction of positive cells compared to vehicle control. ATP-stimulated ppERK levels were determined by subtracting the baseline proportion of positive cells (1 minute of ATP treatment) from the proportion of positive cells after 30 minutes of ATP treatment for each condition: vehicle and 100 nM concentration of each inhibitor. Percent inhibition was determined by taking the ratio of ATP-stimulated ppERK for each drug (at 100 nM) to ATP-stimulated ppERK for vehicle control. Estimates of the mean and error of percentages were determined by sampling the mean proportion of positive cells in 1000 randomly selected cells 10 times.

To quantify the ppERK staining intensity per cell, cells without detectable ppERK staining were assigned a value of –0.5. Average ppERK staining intensity was determined by dividing the integrated signal intensity of ppERK by total cell area and represented as log-transformed values. A ppERK intensity score was calculated using an empirical formula that takes into account the proportion of positive cells and the intensity values of all ppERK-positive cells. The score calculated for the earliest treatment time (1 min) was used as the reference value and the normalized scores represent the ratio of the ppERK scores at each time point over the 1-min time point. For ionomycin treatment time series, the mean values of integrated ppERK intensity per cell per condition are calculated as the mean values from all replicate wells per condition.

#### Fura-2 Spike detection

Automatic classification of cells exhibit calcium spiking behavior from Fura-2 images of BRAFi time series (5612 unique cell traces) was determined using the following criteria: Traces without spikes were identified using all of the following criteria: A) Traces with zero slopes when fit to linear model; B) Traces with mean values <0.05 and standard deviations <0.01; C) Application of DBscan clustering (epsilon=2, minimum number of points=20) to principal component analysis (PCA) of 22 <u>c</u>anonical <u>t</u>ime series <u>ch</u>aracteristics (Catch-22) identified two clusters, one with significant overlap with traces from *a* and *b*, which was assumed to demarcate cells that do not exhibit calcium spikes; D) No values of Fast Fourier Transform values > 10 (indicative of repeating patterns; oscillations); and E) No values above threshold (0.17).

These criteria were able to rapidly classify all 5612 observed Fura-2 ratio traces into spiking versus non-spiking behaviors; however, a significant number of traces classified as exhibiting spikes, when visually assessed for spiking patterns, would not have been considered spiking. Therefore, the estimates of proportions of spiking cells is likely an overestimation of the true proportion of spiking cells. This overestimation appears to be approximately 4–5% based on the automatic detection of cells not treated with BRAFi, which have <1% spiking cells when assessed by visual examination (not shown).

#### Calbryte-520 Spike detection

The automated classification of spiking activity used for Fura-2 could not be applied to Calbryte image data due to a diminishing signal over time and automatic focus correction that resulted in uneven measurements. A subset of 100 randomly selected cells per condition (except where noted) were manually annotated for spiking behavior. The presence of at least one clear spike in Calbryte intensity during the imaging period represented a spiking cell. To rigorously estimate the proportions of spiking cells, bootstrapping was performed on these annotated cells (described below). Under conditions of ATP stimulation, only spikes occurring after the first 10 images were considered to avoid artifacts, but all cells (rather than a random sample) were manually annotated for spiking activity.

#### Bootstrapping to estimate mean proportions of spiking cells for each condition

Confidence intervals for the proportion of spiking cells in a population were determined using bootstrapping based on the number of cells for which spiking annotation was obtained. Sampling of at least 20% of the annotated cells (minimum of 50 cells per condition) was performed at least 10 times per condition. For example, since all cells in the data from Fura-2 imaging were annotated (∼900–1200 cells per condition), proportions of positive cells were calculated from 200 randomly sampled cells (with replacement) 100 times. For Calbryte images, fewer cells were annotated for spiking activity (typically between 160 and 500 cells per condition.) For these experiments, 50 randomly sampled cells (with replacement) were used to calculate the percentage of spiking cells 100 times for each condition.

**Supplementary Table 1.** Reagent information.

| Compound | Solvent | Vendor |
| --- | --- | --- |
| PLX4720 | DMSO | MedChemExpress, HY-51424 |
| Dabrafenib | DMSO | MedChemExpress, HY-14660 |
| Trametinib | DMSO | MedChemExpress, HY-10999 |
| AACBA hydrochloride | MilliQ water | Alamone Labs, A-410 |
| AZD9056 hydrochloride | DMSO | MedChemExpress, HY-19427A |
| A740003 | DMSO | MedChemExpress, HY-50697 |
| BX430 | DMSO | Selleck Chem, S0758 |
| isoPPADS tetrasodium salt | MilliQ water | Tocris, 0683 |
| Gefapixant | DMSO | Selleck Chem, S6664 |

## ACKNOWLEDGEMENTS

We would like to acknowledge the Vanderbilt High Throughput Screening facility for their contributions to this work and that some results published here are in whole or part based upon data generated by the TCGA Research Network: https://www.cancer.gov/tcga. Funding was provided by U54CA217450 to VQ; R50CA243783 to DRT; T32MH064913 to PES; and DK129340 and DK136768 to DAJ.

