## Supplementary Figures for "Purinergic Ca^2+^ signaling as a novel mechanism of drug tolerance in BRAF mutant melanoma"

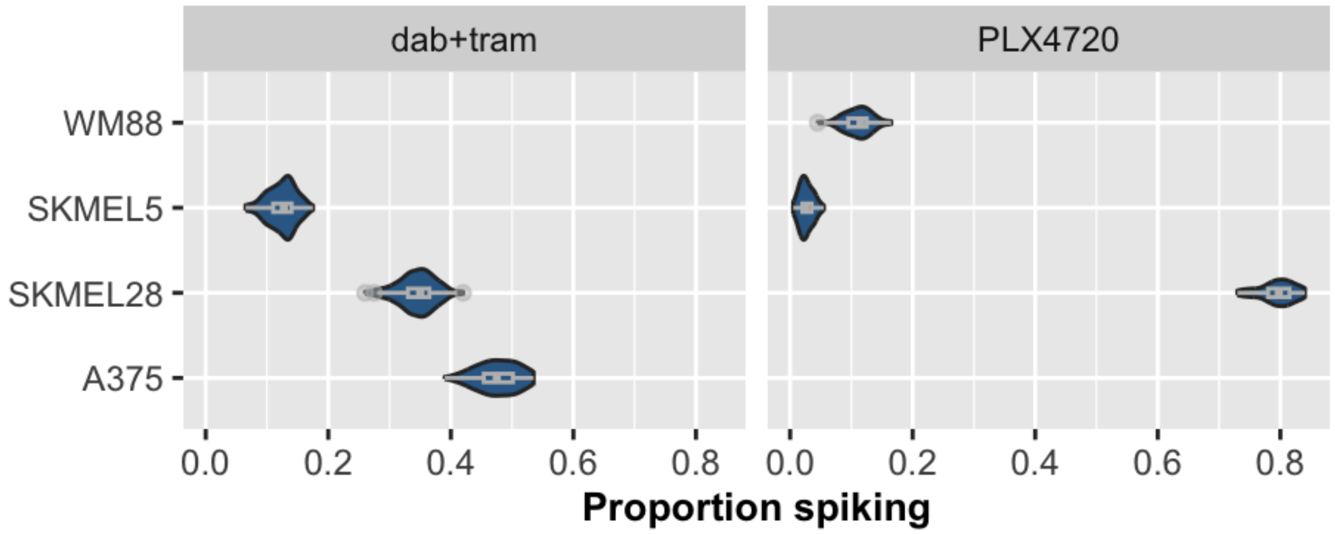


Figure S1. **Spiking occurs in numerous BRAF mutant melanoma lines and under different conditions of treatment.** Indicated cell lines were treated for 3 days with either 8 µM PLX4720 or 1 µM dabrafenib + 100 nM trametinib (dab + tra) and imaged in serum containing Imaging Buffer (+10% FBS). Traces were manually analyzed for spiking activity. Bootstrapping was performed to generate proportions of spiking cells presented in the figure.

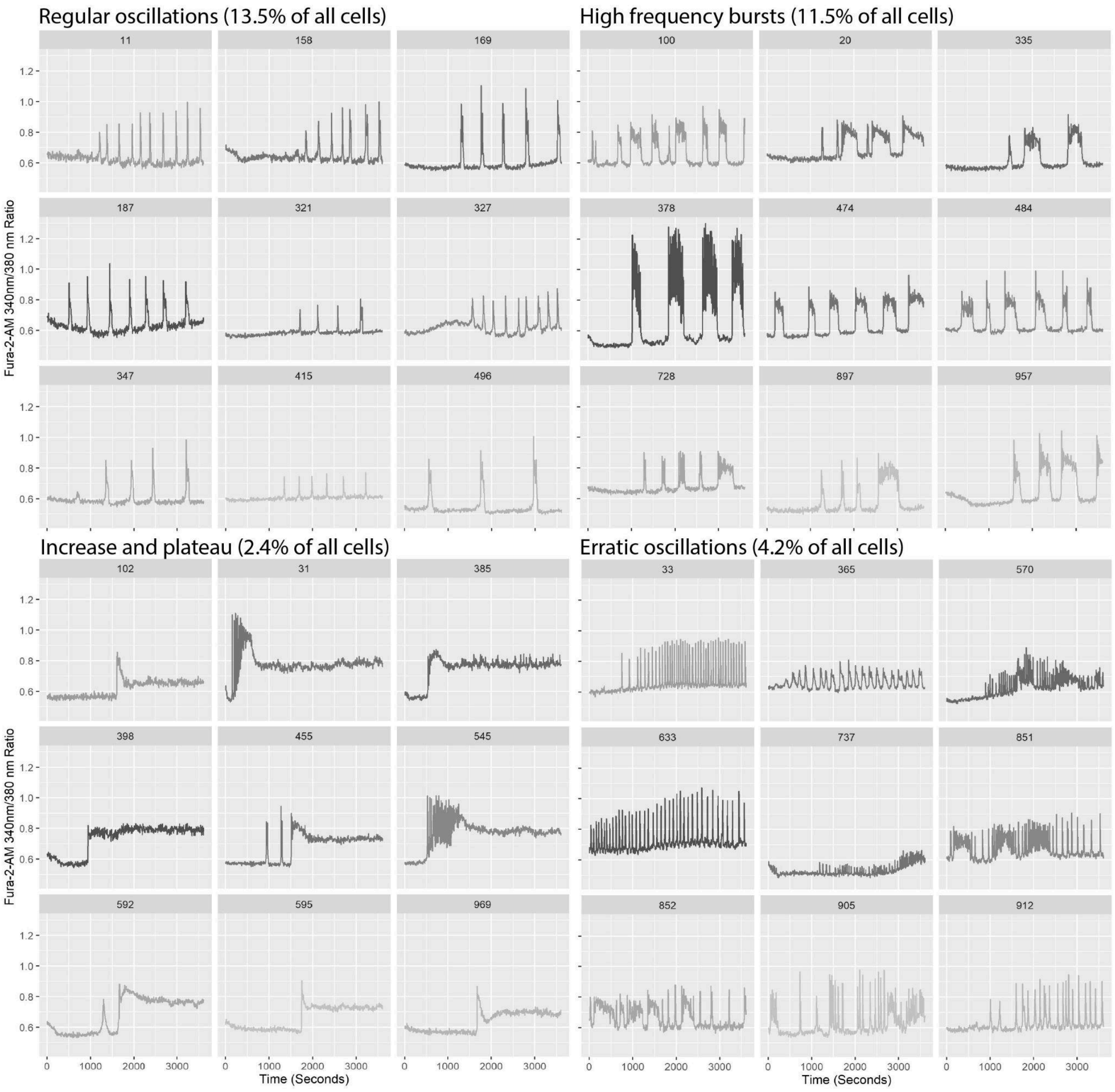


Figure S2. **Spiking activity can be qualitatively grouped by spike pattern.** Fura-2 $Ca^{2+}$ imaging (60 minutes) of A375 cells treated with 8 µM PLX4720 for 14 days. Four spike patterns became apparent upon close examination and are represented above. All pulsing cells with more than two spikes were manually gated into one of these four groups. Notably, some traces have mixes of spike patterns (i.e. high frequency bursts may repeat at regular frequencies and/or have single spikes interspersed). Cells with only one or two spikes were not grouped in the above and accounted for approximately 38% of the spiking population. Traces presented here are representative of the population, to the best of our ability.

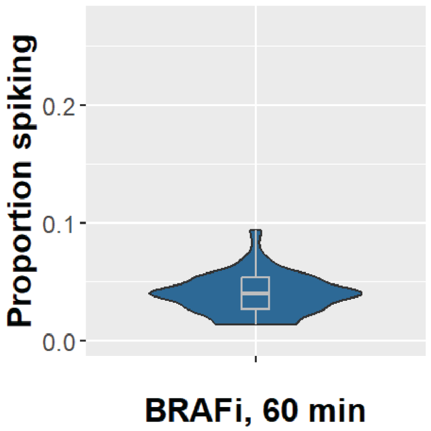


Figure S3. **Spiking activity in drug-sensitive cells is minimal.** Proportion of spiking A375 cells after 60 minutes of BRAFi is approximately the same proportion found at 0 days BRAFi in Fig. 1C. Data was acquired using calbryte-520. Traces were manually analyzed for spiking activity. Bootstrapping was performed to generate proportions of spiking cells presented in the figure.

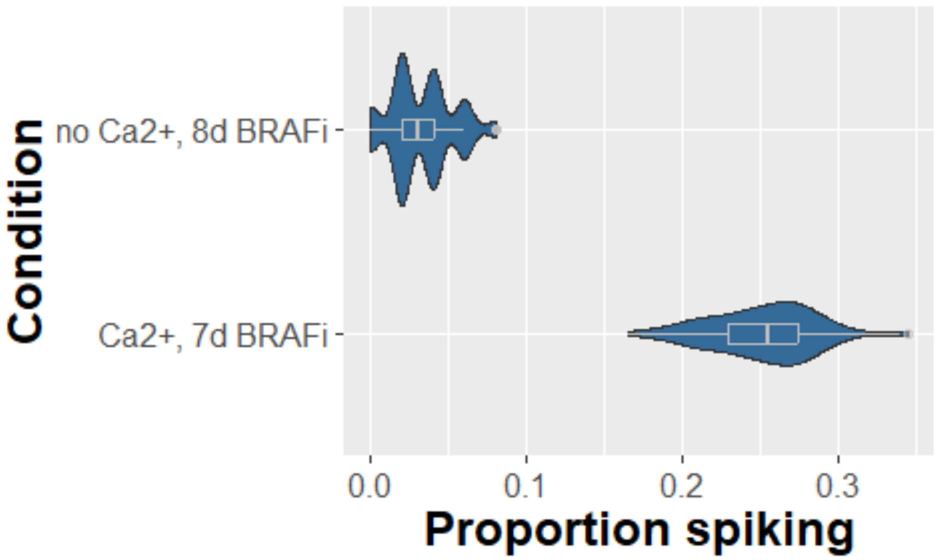


Figure S4. **Spiking activity in drug-tolerant cells treated for 8 days relies on extracellular $Ca^{2+}$.** A375 cells were treated with 8 µM PLX4720 for indicated periods of time and imaged with Fura-2 for 40 minutes. 7 days of BRAFi treatment were taken from Fig. 1C for comparison to 8 days BRAFi without $Ca^{2+}$; since 10 days BRAFi has a similar proportion spiking as 7 days (Fig. 1C), we consider comparing the above conditions is appropriate. $Ca^{2+}$-free HBSS was used as the imaging and wash buffer for the no $Ca^{2+}$ condition.

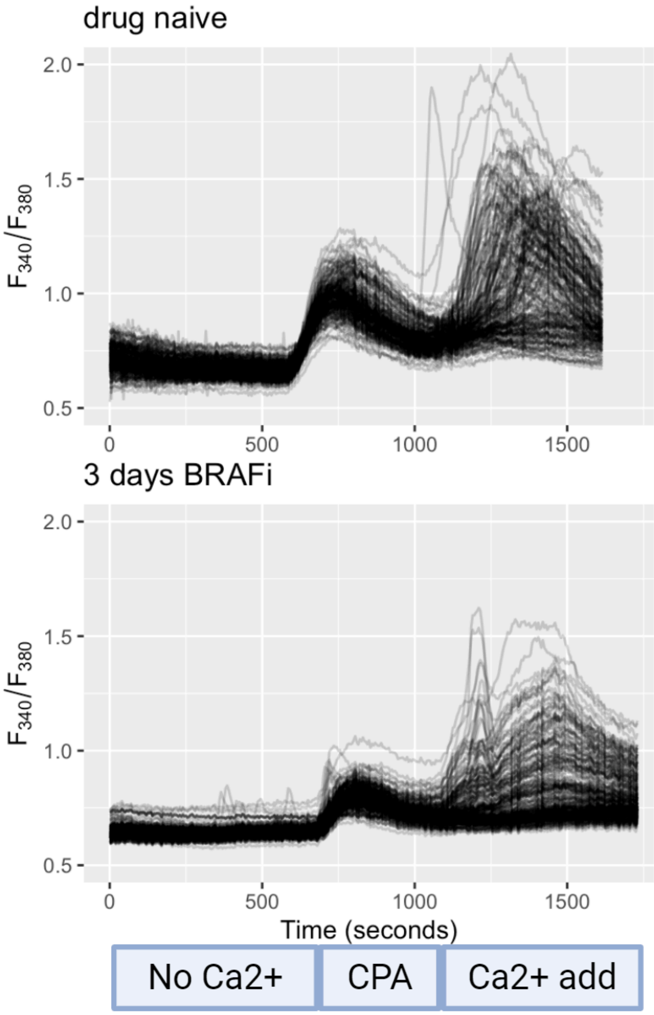


Figure S5. **Single cell $Ca^{2+}$ traces in store operated $Ca^{2+}$ entry (SOCE) assays performed on BRAFi naive and 3 day BRAFi treated cells**. Cells were loaded with the $Ca^{2+}$ dye Fura-2-AM and equilibrated in $Ca^{2+}$-free buffer before the SERCA inhibitor, cyclopiazonic acid (CPA), was added to release free ER $Ca^{2+}$ into the cytoplasm. Quantification of ER $Ca^{2+}$ levels occurs during the phase labeled "CPA". Seven minutes after CPA treatment, extracellular $Ca^{2+}$ is added to measure SOCE activity. Quantification of SOCE activity levels occurs during the phase labeled "Ca2+ add". Cytoplasmic $Ca^{2+}$ clearance can also be calculated from the single-cell traces during the "CPA" phase.

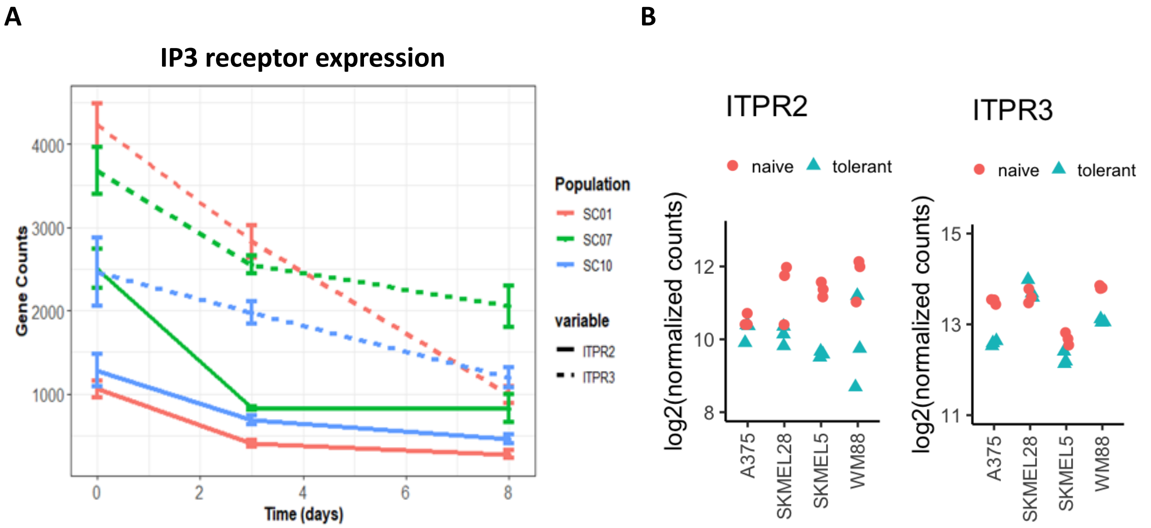


Figure S6. **IP3 receptor expression in BRAFi.** *A*) Data from SKMEL5 subclones (SC01, SC07, SC10) treated with 8 µM BRAFi for the indicated times. *B*) Transcript abundance from drug-naive and drug-tolerant (8 days in 8 µM BRAFi) melanoma cells.

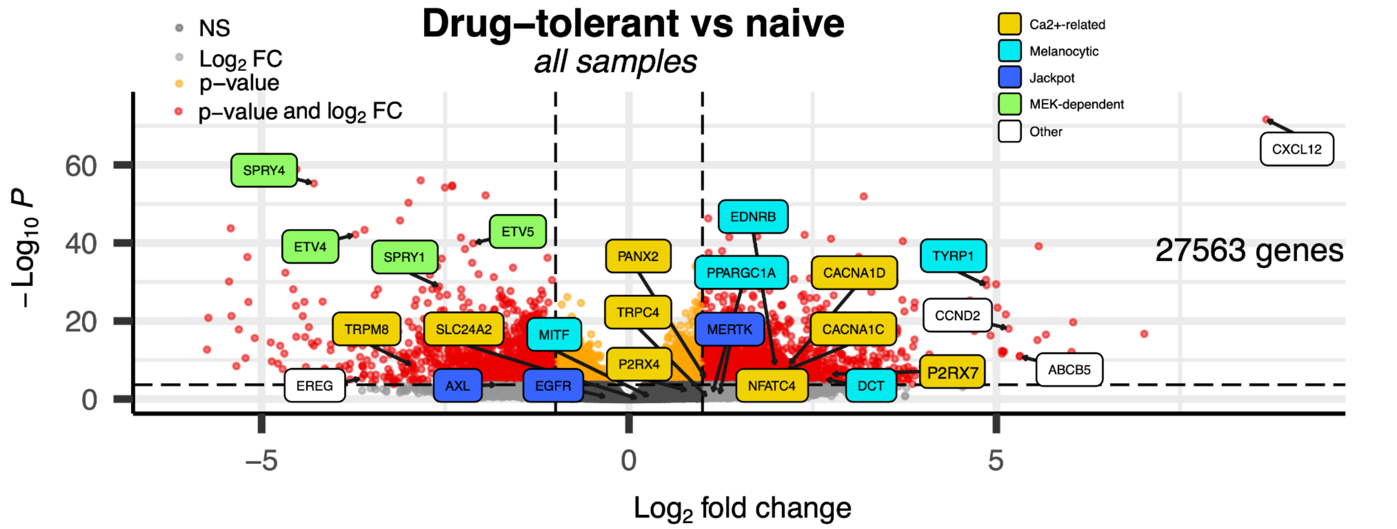


Figure S7. **Volcano plot of up- and down-regulated genes in drug-tolerant versus BRAFi-naive melanoma cells.** Melanoma cells were treated with 8 µM BRAFi for 8 days (drug-tolerant) or left untreated (naive). Whole transcriptome analysis (bulk RNAseq) was performed on four BRAF mutant melanoma cell lines and analyzed together. Colors indicate distinct classes of genes, related to $Ca^{2+}$ transport and signaling, melanocyte differentiation, Jackpot cells, and a signature of MEK-dependent signaling.

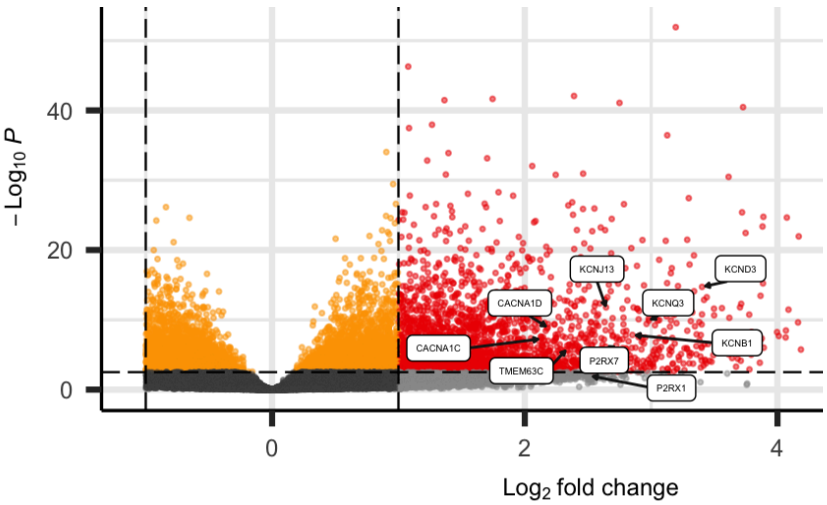


Figure S8. **Volcano plot of up- and down-regulated genes in melanoma cells under BRAFi conditions** for 8 days, a treatment previously shown to induce an idling population of drug-tolerant cells. Data are from bulk RNAseq of four BRAF mutant melanoma cell lines that were tested. Highlighted are genes related to $Ca^{2+}$ and monoatomic cation channels (GO 5261).

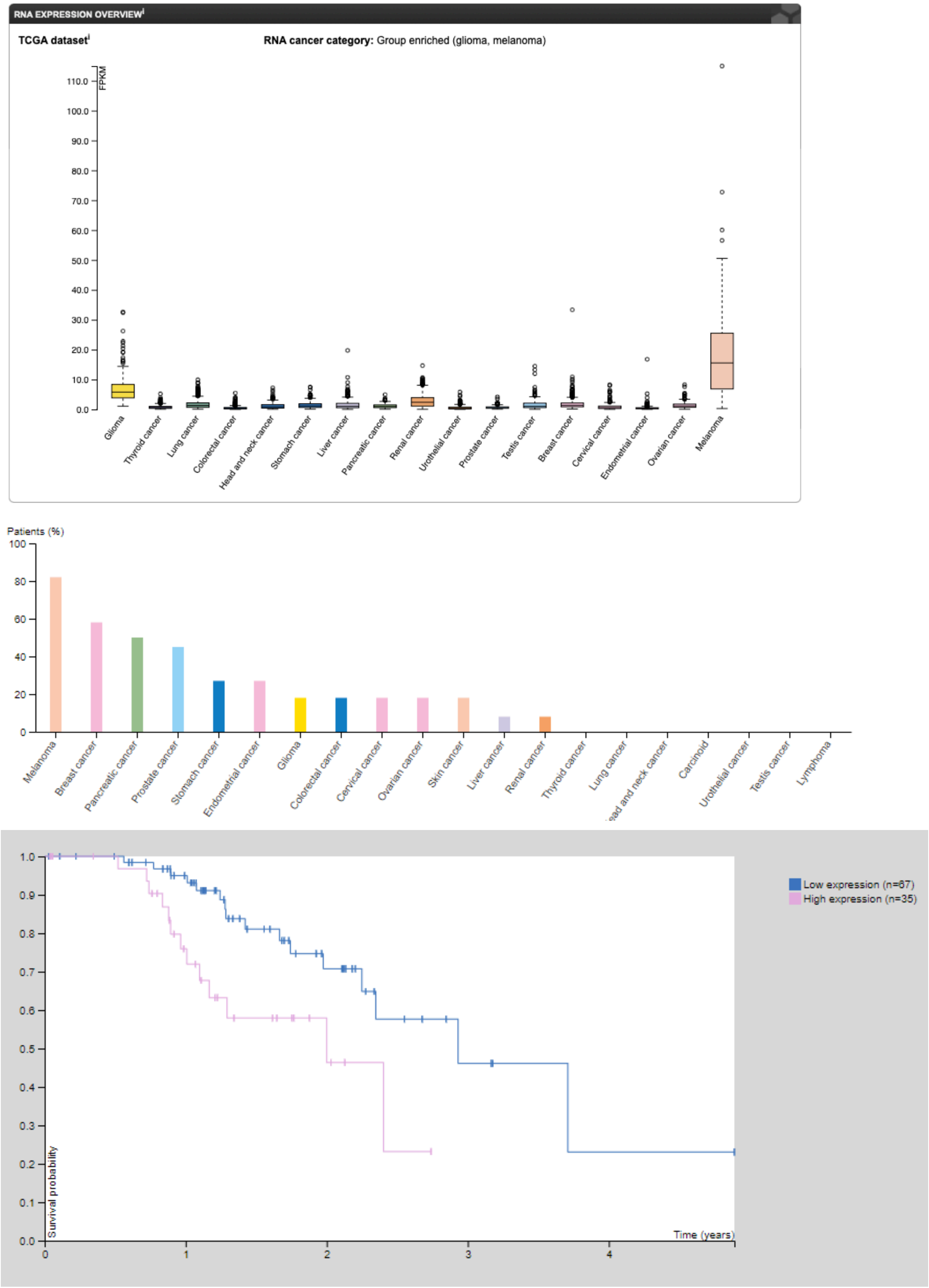


Figure S9. **TCGA data demonstrating high levels of P2RX7 expression in melanoma.** Figures generated by the Human Protein Atlas. *Top* mRNA expression levels in different cancer types. *Middle* Antibody (HPA034968) detection of P2X7 in patients with different cancers. 9 out of 11 patients showed medium to high expression, and all 11 patients showed at least low levels of expression. *Bottom* Survivorship curves of patients with high or low mRNA expression as determined by a cutoff value of 23.34 FPKM.

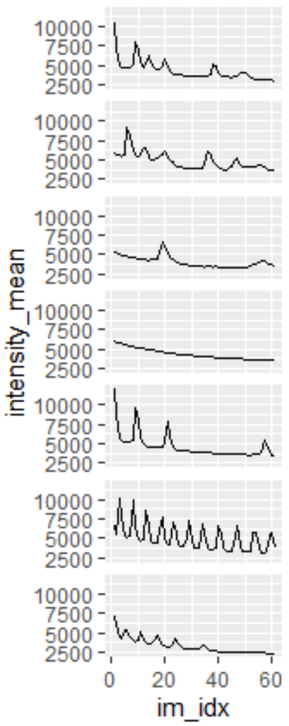


Figure S10. **ATP stimulation generates spikes of $Ca^{2+}$ in drug tolerant cells.** Representative traces from ATP stimulated (39 μM) $Ca^{2+}$ imaging in drug-tolerant A375 cells. Data traces are recorded over 5 minutes with an image recorded every 5 seconds. These cells were selected to be representative of those observed to experience obvious $Ca^{2+}$ spikes.

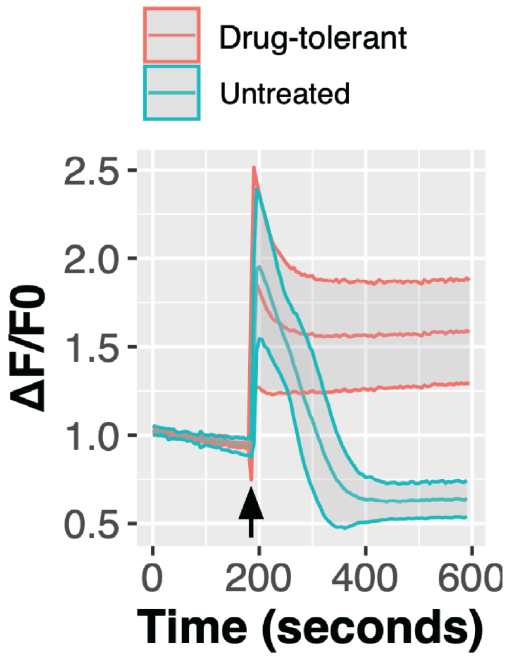


Figure S11. **ATP stimulation of drug-tolerant idling cells.** A375 cells treated with 8 µM PLX4720, or vehicle, for 8 days were imaged with Fura2. A background was recorded for 3 minutes before 10 mM ATP was added on top of cells (arrow). Average $Ca^{2+}$ trace with 95% confidence intervals are displayed for all cells imaged.

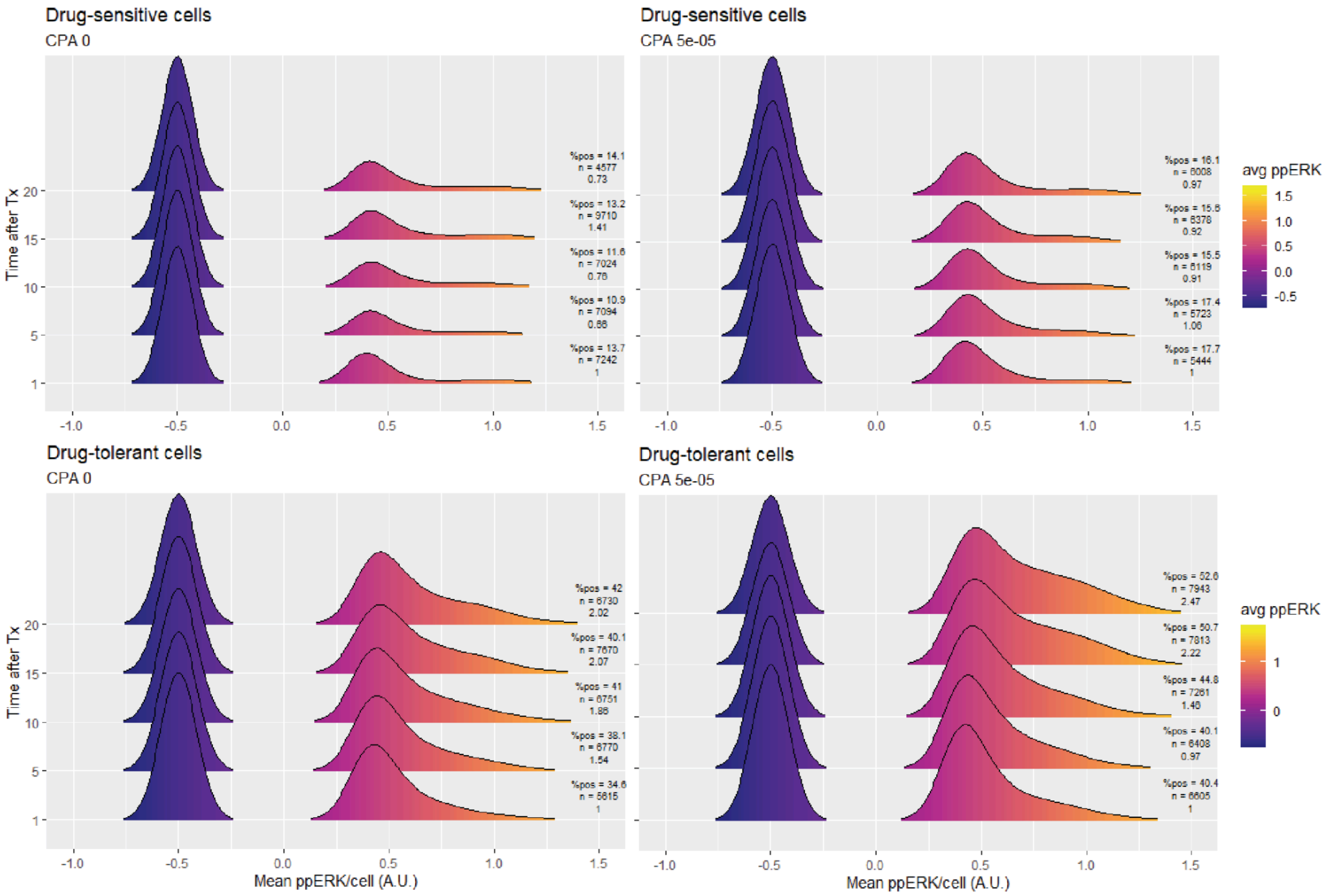


Figure S12. **CPA induces ERK phosphorylation in drug-tolerant, but not drug-sensitive cells.** Drug-sensitive (60 min BRAFi) *(top)* and drug-tolerant (3 days BRAFi) *(bottom)* A375 cells were treated with 50 µM CPA to induce a cytoplasmic $Ca^{2+}$ signal. Cells were incubated in CPA, fixed with 4% PFA, and stained for ppERK. Cells without detectable ppERK staining were given a value of –0.5 and the minimum ppERK level in the positive cells was set to 0. Drug-sensitive cells displayed little to no ppERK activation, whereas drug-tolerant cells did, as expected.

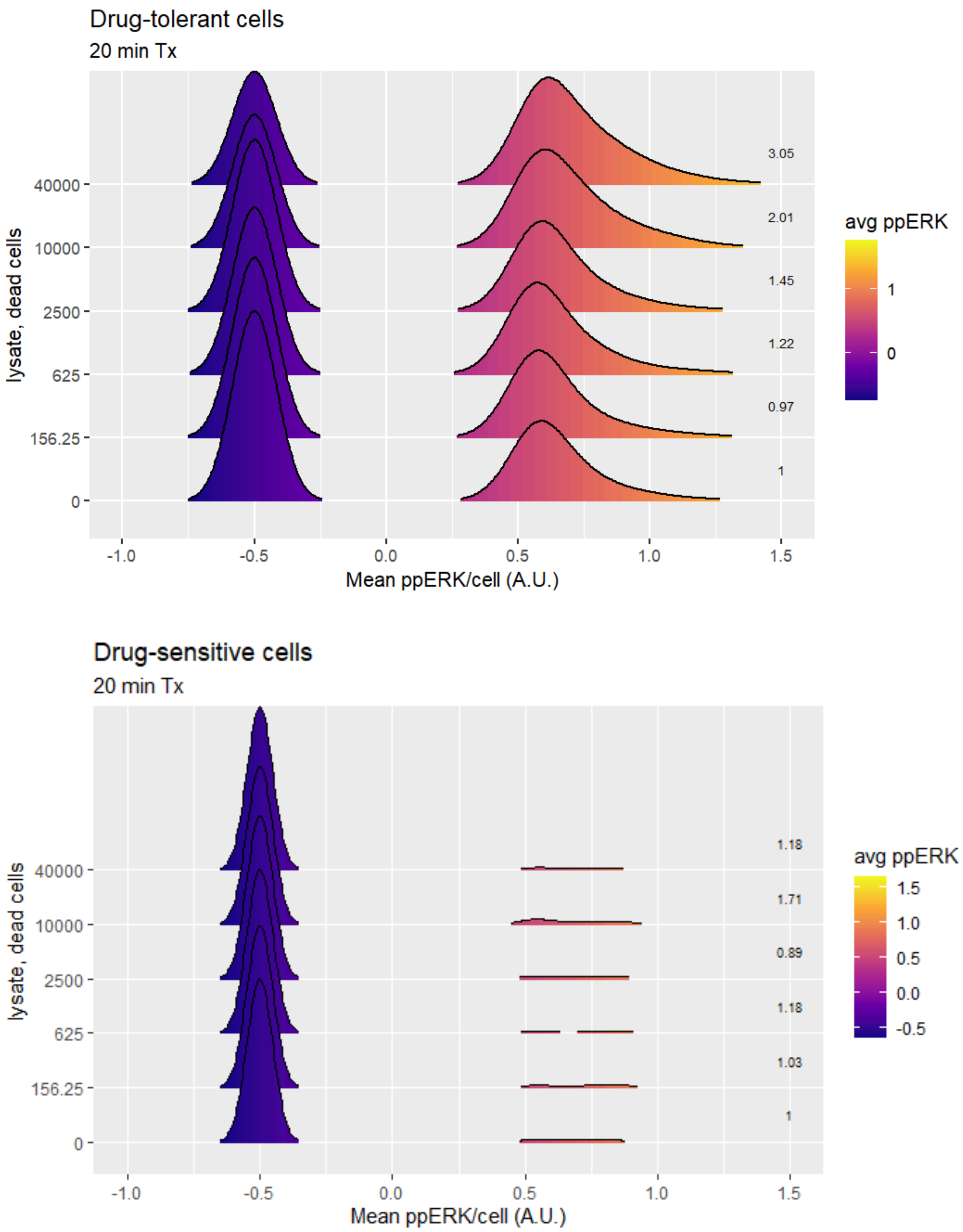


Figure S13. **Cell lysate stimulates ppERK in drug-tolerant but not drug sensitive cells.** Drug naive cells were trypsinized, washed, and boiled at 98°C for 10 minutes, and centrifuged at 20,000xg for 5 minutes. Supernatant were diluted and incubated on drug-tolerant (3 days BRAFi) or drug-sensitive (60 min BRAFi) cells for 20 minutes before fixation and staining for ppERK. Quantification of the staining intensity (normalized to 0) are shown on the right, on each bipartite density distribution.

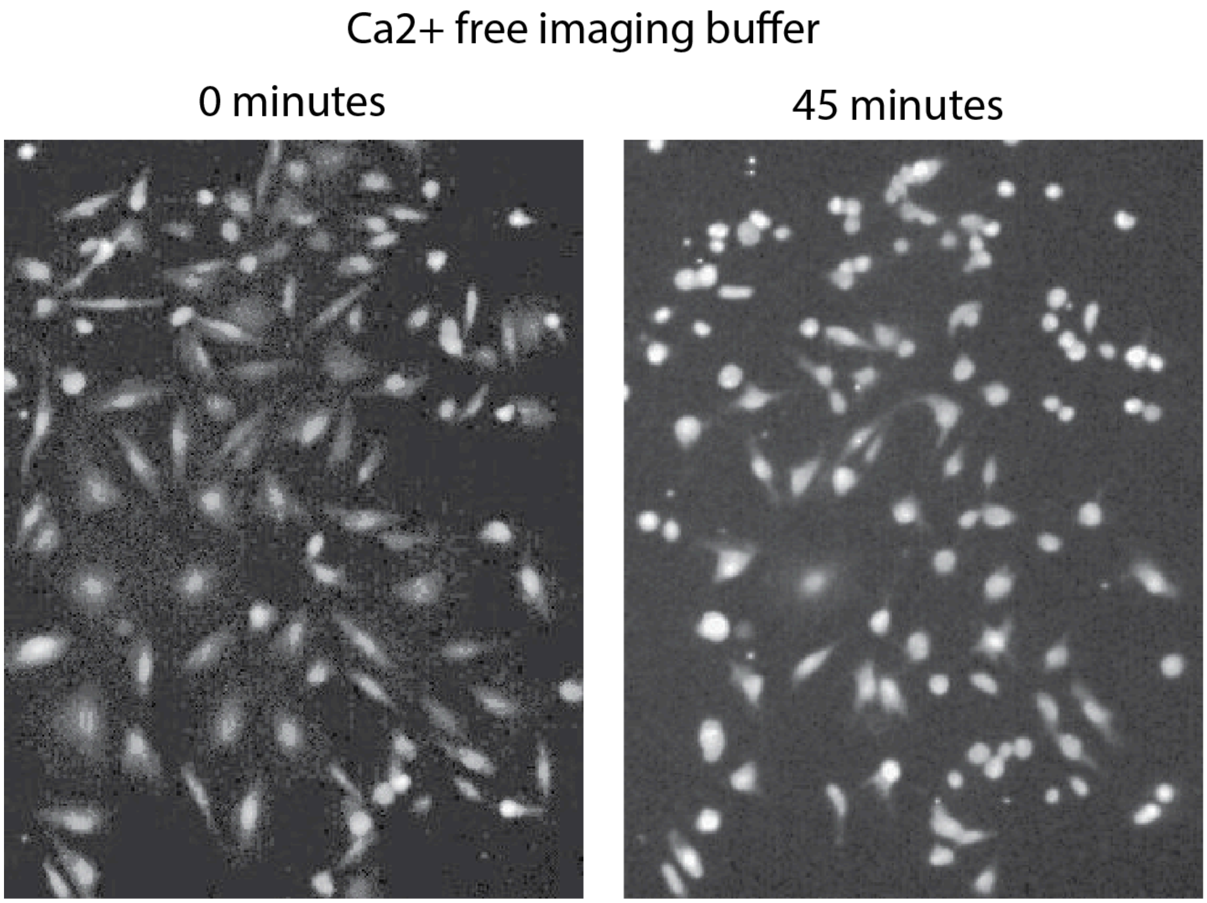


Figure S14. **Drug-tolerant cells do not tolerate $Ca^{2+}$ free conditions.** *(Left)* Drug-tolerant A375 cells (8 days BRAFi) immediately after being switched to $Ca^{2+}$ free imaging buffer (HBSS). *(Right)* Those same cells after 40 minutes in $Ca^{2+}$ free HBSS. Images were captured using Fura2 and are representative of the population. Cells do not ball up when imaged in $Ca^{2+}$:HBSS, suggesting this effect is not related to imaging and/or phototoxicity.

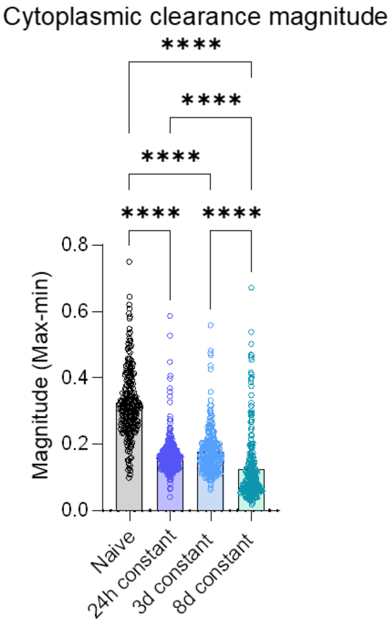


Figure S15. **Quantification of single cell SOCE $Ca^{2+}$ traces.** Data represents quantification of max $Ca^{2+}$ levels subtracted from lowest during ER release phase. A375 cells were treated for indicated periods of time with 8 μM PLX4720.
